# FAM76B regulates NF-κB-mediated inflammatory pathway by influencing the translocation of hnRNPA2B1

**DOI:** 10.1101/2022.12.29.522198

**Authors:** Dongyang Wang, Xiaojing Zheng, Lihong Chai, Junli Zhao, Jiuling Zhu, Yanqing Li, Peiyan Yang, Qinwen Mao, Haibin Xia

## Abstract

FAM76B has been reported to be a nuclear speckle localized protein with unknown function. In this study, FAM76B was first demonstrated to inhibit the NF-κB-mediated inflammatory pathway by affecting the translocation of hnRNPA2B1 *in vitro.* We further showed that FAM76B suppressed inflammation by regulating the NF-κB pathway *in vivo* using a traumatic brain injury (TBI) model in FAM76B knockout mice. Lastly, FAM76B was shown to interact with hnRNPA2B1 in human tissues taken from patients with acute, organizing, and chronic TBI, and with different neurodegenerative diseases. The results suggested that FAM76B mediates neuroinflammation by influencing the translocation of hnRNPA2B1 *in vivo* during TBI repair and neurodegenerative diseases. In summary, we for the first time demonstrated the role of FAM76B in regulating inflammation and further showed that FAM76B could regulate the NF-κB-mediated inflammatory pathway by affecting hnRNPA2B1 translocation, which provides new information for studying the mechanism of inflammation regulation.

## 1. Introduction

Peripheral immune cells mediating inflammation have been reported to be closely associated with the development of some diseases, such as cancer (Moore et al. 2010; Kay et al. 2019; Suarez-Carmona et al. 2017; Khansari et al. 2009; Khandia et al. 2020; Kolb et al. 2016), obesity (Kawai et al. 2021; Khodabandehloo et al. 2016; Seong et al. 2019; Saltiel et al. 2017; Curley et al. 2021), and autoimmune diseases (Kumar. 2019; Xie et al. 2019; Lochhead et al. 2021; Venkatesha et al. 2016; Abou-Raya et al. 2006), among others. Microglia, as the resident macrophages of the central nervous system (CNS), act as the first line of defense in the brain (Muzio et al. 2021) and play a role in mediating neuroinflammation. It has been demonstrated that neuroinflammation is an important characteristic of almost all neurological disorders (Gilhus et al. 2019; Brambilla. 2019), which is the common thread that connects brain injuries to neurodegenerative diseases, and researchers have provided evidence of this commonality using traumatic brain injury (TBI) as an example. Inflammation is regulated by different signaling pathways (Kumar et al. 2003; Kim et al. 2004; Rawlings et al. 2004; Hawkins et al. 2015; Lawrence. 2009); however, the detailed mechanisms regulating inflammation are still poorly understood.

Human FAM76B is a 39 kDa nuclear speckle-localized protein that consists of 339 amino acids. It contains homopolymeric histidine tracts that are considered a targeting signal for nuclear speckles (Alvarez et al. 2003; Herrmann et al. 2001; Salichs et al. 2009). Although the function of FAM76B is still unknown, many poly (His)-containing proteins have been shown to be involved in DNA- and RNA-related functions and are overrepresented during development of the nervous system (Salichs et al. 2009). In our previous study, by using immunohistochemical staining with custom-made anti-hFAM76B monoclonal antibodies (Zheng et al. 2016), we found strong immunolabeling of FAM76B in the human brain, lymph nodes, and spleen. The results raised the question: does this protein play a role in regulating inflammation and neuroinflammation? In this study, we for the first time demonstrate that FAM76B can inhibit inflammation *in vitro* and *in vivo* by regulating the NF-κB pathway. Furthermore, we showed that FAM76B can regulate the NF-κB pathway and mediate inflammation by affecting the translocation of hnRNPA2B1.

## 2. Materials and Methods

### Real-time PCR for gene expression in tissues and cell lines

Total cellular RNA was isolated from cells by TRIzol (Invitrogen, Carlsbad, CA, USA). The cDNA was synthesized using a PrimeScript^TM^ RT reagent Kit (Takara Bio, Beijing, China) according to the manufacturer’s manual. Gene expression was performed using a real-time PCR kit (Thermo, Rockford, IL, USA) and was normalized to GAPDH. The primers used are listed in Table S1.

### Generation of FAM76B knockdown and knockout U937 cell lines by lentivirus-mediated Cas9/sgRNA genome editing

An efficient sgRNA (GCAGGGACTGTGGAAACAG) targeting Exon 6 of the human FAM76B gene was screened using a T7E1 assay. The primers used for the experiment are listed in Table S2. To generate U937 cell lines with FAM76B knockdown, an inducible lentiviral vector was constructed. Briefly, the Bi-Tet-On inducible system, previously established by our lab (Chen et al. 2015), and a U6-sgRNA expression cassette were introduced into the lentiviral vector pCDH-CMV-MCS-EF1-Puro to generate the inducible lentiviral vector pCDH-Tet-On cassette-U6 sgRNA cassette-EF1-puro. Then, Cas9 and the corresponding sgRNAs were cloned into this lentiviral vector to generate the final vector, pCDH-Tet-on-Cas9-U6-FAM76B sgRNA-EF1-puro. The lentivirus was produced in HEK293T cells. U937 cells were infected by the above lentivirus, screened by puromycin, and then cultured with 2 μg/ml doxycycline for six days. The fresh medium containing doxycycline was changed every three days to induce expression of Cas9. Then, FAM76B knockdown U937 cells were obtained and evaluated by T7E1 assay. Through several rounds of cell cloning by serial dilution of the FAM76B knockdown U937 cells, the FAM76B knockout U937 cell line was obtained and confirmed by gene sequencing and Western blot. In addition, U937 cells were infected by the lentivirus LV-Tet-on-Cas9-EF1-puro to obtain the control cell line.

### Mice

All animal studies were performed in accordance with institutional guidelines and with approval by the Institutional Animal Care and Use Committee of Shaanxi Normal University. C57BL/6 mice were obtained from the animal center of Shaanxi Normal University. The mice were maintained in a controlled environment (12/12 h light/dark cycle, 23±1°C, 55±10% humidity) and given free access to food and water.

### Generation of FAM76B gene trap mutant mice and genotyping

Homozygous FAM76B knockout (*Fam76b^-/-^*) mice were produced using a commercial service, Texas A&M Institute for Genomic Medicine (College Station, TX, USA), by gene trap mutagenesis techniques. Two germline-competent male chimeras were generated and bred with C57BL/6 female mice. To identify the embryos homozygous for the gene trap insertion, PCR was performed on cDNA from mouse embryonic fibroblasts derived from embryonic day 13.5 (E13.5) embryos, using mouse *Fam76b* primers (Table S2). The genotypes of the offspring were determined using tail clip genomic DNA. To detect the wild-type (WT) Fam76b allele, PCR was performed using Fam76b forward and reverse primers (Table S2). To detect the homozygotes, PCR was performed using Fam76b forward and V76 reverse primers (Table S2). The expected size of the fragment from the WT allele was 494 bp. The expected size of the product from intron 1 of the gene trap-containing Fam76b gene was 354 bp.

### Isolation of bone marrow-derived macrophages (BMDMs)

BMDMs were isolated from *Fam76b^-/-^* and WT C57BL/6 mice by flushing the femur with Dulbecco’s modified Eagle medium (DMEM) supplemented with 3% fetal bovine serum (FBS). Cells were plated for 4 h to allow the non-monocytes to adhere to the surface. Then, the monocytes in the culture supernatant were collected and centrifuged at 250 × g and then seeded in DMEM supplemented with 10% FBS, 1% penicillin/streptomycin, 1% L-glutamate, and 10 ng/ml macrophage colony-stimulating factor (Sino Biological Inc., Beijing, China). The fresh medium containing 10 ng/ml macrophage colony-stimulating factor was changed every two days to induce differentiation of the cells into macrophages. After six days in culture, BMDMs were collected for the following experiments.

### Isolation of mouse embryonic fibroblasts

Embryos from WT and *Fam76b^-/-^* mice were isolated at about E13.5. After the heads, tails, limbs, and most of the internal organs were removed, the embryos were minced and digested in 0.1% (m/v) trypsin for 30 minutes at 37°C and then centrifuged at 250 × g to pellet mouse embryonic fibroblasts. The mouse embryonic fibroblasts were cultured in DMEM supplemented with 10% FBS, 1% penicillin/streptomycin, and 1% L-glutamate.

### Lipopolysaccharide (LPS) treatment of mice

For brain injection, LPS (Sigma-Aldrich, St. Louis, Missouri, USA) (2 μg/μl, 1.5 μl) was injected into the prefrontal cortex using a microsyringe at the following stereotaxic coordinates: 2.60 mm caudal to bregma, 2 mm lateral to the midline, and 1.7 mm ventral to the surface of the dura mater. Vehicle (0.5% methylene blue in phosphate-buffered saline [PBS] or PBS only) was injected in a similar manner into the contralateral prefrontal cortex. For intraperitoneal injection, mice were injected intraperitoneally with LPS (5 μg/g body weight) or with PBS once a day for two days before the spleens were harvested for pathologic evaluation.

### Rescue of FAM76B in FAM76B knockout U937 cells or BMDMs

For FAM76B knockout U937 cells, cells were infected by the lentivirus LV-CMV-hFAM76B-EF1-GFP or the control lentivirus LV-CMV-MCS-EF1-GFP. Three days after infection, cells were prepared for LPS treatment. For BMDMs from FAM76B knockout mice, cells were infected by the lentivirus LV-CMV-mFAM76B-EF1-GFP or the control lentivirus LV-CMV-MCS-EF1-GFP. Six days after infection, cells were prepared for LPS treatment.

### LPS treatment of cells

To evaluate the effects of FAM76B on the cytokine production of U937 cells or BMDMs, different U937 cell lines or BMDMs from FAM76B knockout mice were treated with LPS. Briefly, for the impact of FAM76B on cytokine production, FAM76B knockdown U937 cells were plated in 6-well plates (5×10^5^ cells/well) and treated with 10 ng/ml PMA (Sigma-Aldrich, St. Louis, Missouri, USA) in association with 10 ng/ml LPS and 20 ng/ml hIFNγ (Sino Biological Inc., Beijing, China) for 48 h, while FAM76B^-/-^ U937 cells were plated in 6-well plates (5×10^5^ cells/well) and treated with 1ng/ml PMA for 48 h, and then were treated with 10 ng/ml LPS and 20 ng/ml hIFNγ for 24 h. For rescuing FAM76B in the FAM76B^-/-^ U937 cell line, U937 cells were plated in 6-well plates (5×10^5^ cells/well) and treated with 0.5 ng/mL PMA for 24 h and then stimulated with 1 ng/mL LPS and 20 ng/ml hIFNγ for 24 h. For rescuing FAM76B in FAM76B^-/-^ BMDMs, BMDMs were plated in 6-well plates (2×10^5^ cells/well) and treated with 1 ng/mL LPS and 20 ng/mL mIFNγ (Sino Biological Inc., Beijing, China) for 24 h. After treatment, cells were harvested in TRIzol for the detection of cytokine expression by real-time PCR.

### IL-6 promoter activity assay

Human IL-6 promoter (1868 bp) was obtained by PCR using HEK293 cells genomic DNA as PCR template. The IL-6 promoters were confirmed by gene sequencing and then were inserted into pGL3 basic plasmid to obtain the human IL-6 promoter activity reporter vector pGL3-IL6-Luc. Human P50 and P65 cDNA was obtained by PCR using the MegaMan Human Transcriptome Library as the template. Human P65 and P50 were confirmed by gene sequencing and then were inserted into the eukaryotic expression vector pAd5 E1-CMV-MCS to obtain the human P65 and P50 expression vectors pAd5 E1-CMV-P65 and pAd5 E1-CMV-P50, respectively. WT HEK293 cells and FAM76B^-/-^ HEK293 cells were plated into 24-well plates at a density of 2×10^5^ cells per well. The next day, the promoter activity reporter vector pGL3-IL6-Luc (200 ng) and the Renilla luciferase expression vector pRL-CMV (50 ng) as the internal reference plasmid were co-transfected with the P65 and P50 expression vector pAd5 E1-CMV-P65/P50 (each 300 ng) or the control vector pAd5 E1-CMV-MCS (600 ng) into the cells using X-tremeGENE HP Reagent (Roche, Indianapolis, IN, USA) according to the manufacturer’s protocol. 48 h later, the cells were collected for a luciferase activity assay using a dual-luciferase assay kit (Promega, Madison, WI, USA). The normalized luciferase activity was obtained by using the formula: Normalized luciferase value = Fly luciferase value/Renilla luciferase value.

Meanwhile, the human IL-6 promoter was inserted into the upstream of luciferase in the lentivirus vector pCDH-luciferase-EF1-Neomycin to obtain the vector pCDH-IL-6 promoter-luciferase-EF1-Neomycin. Then, the WT or FAM76B^-/-^ U937 cells were infected by the above lentivirus. After selected by G418, WT and FAM76B^-/-^ U937 cells were seeded into 24 wells (2×10^5^ cells per well) and incubated with different concentrations of LPS for 36 h. The cells were then collected for a luciferase activity assay.

Two kinds of human NF-κB binding motif (motif 1: GGGAATTTCC, motif 2: GGGATTTTCC) were also synthesized and inserted into the upstream of miniCMV promoter in the lentivirus vector pCDH-miniCMV-luciferase-EF1-Neomycin to obtain the vector pCDH-NF-κB binding motif 1(or 2)-miniCMV-luciferase-EF1-Neomycin. Then, the WT or FAM76B^-/-^ U937 cells were infected by the above two kinds of lentivirus. After selected by G418, WT and FAM76B^-/-^ U937 cells were seeded into 24 wells (2×10^5^ cells per well) and incubated with different concentrations of LPS for 36 h. Then the cells were collected for a luciferase activity assay.

### Flow cytometry

Spleens from five-month-old WT and *Fam76b^-/-^* mice were pressed through an 80 μm mesh with a syringe plunger to acquire single-cell suspensions. Red blood cells were removed from single-cell suspensions using red blood cell lysis buffer (Shanghai Yeasen Biotechnology, Shanghai, China). The cells were then washed twice and resuspended in 200 μl of wash buffer for final flow cytometric analysis. Fluorescent-labeled antibodies used for flow cytometry are listed in Table S3, and flow cytometry staining was performed according to the manufacturer’s instructions using propidium iodide solution (BioLegend, San Diego, CA, USA) to exclude dead cells.

### Western blot

Cells were lysed by RIPA buffer. Cell lysis was subjected to SDS-PAGE and subsequently blotted onto methanol pretreated polyvinylidene difluoride (PVDF) membranes. The PVDF membranes were incubated with the primary antibody overnight at 4°C. Membranes were washed and incubated with corresponding Horseradish Peroxidase (HRP)-conjugated secondary antibody. The membranes were visualized using enhanced chemiluminescence Western blot detection reagents (Thermo Fisher, Waltham, MA, USA) in a chemical luminescence imaging apparatus according to the manufacturer’s protocol. Primary antibodies used for Western blot are listed in Table S3.

### Experimental TBI by controlled cortical impact (CCI)

Experimental TBI was performed using the CCI model, as described (Zheng et al. 2022). Briefly, mice were anesthetized with 1.5% pentobarbital sodium at a dose of 50 mg/kg. The mice were fixed in a stereotactic frame and subjected to a craniotomy (5 mm diameter) in the right parietal region using a motorized drill. The brain was exposed to a pneumatic impactor (Brain injury device TBI-68099, RWD, China), and TBI was produced using the following parameters: diameter impactor tip, 3 mm diameter metal tip; velocity, 3.3 meters/second; duration, 0.1 second; depth of penetration, 2 mm. After the wound was sutured, an electric heater was used to maintain the animals’ body temperature until they were completely awake and able to move freely, which occurred approximately 1–2 h after the injury. Buprenorphine was diluted in 0.9% NaCl to a concentration of 0.01 mg/ml, and a 0.1 mg/kg dose was administered subcutaneously, which provided 72 h of sustained post-operative analgesia. Sham-operated WT and *Fam76b* knockout mice, used as controls, were treated the same as the CCI-treated mice, except for the craniotomy and CCI.

### Cytokine expression of mouse brains by enzyme-linked immunosorbent assay (ELISA) and real-time PCR

The mice were perfused transcardially with ice-cold PBS, and prefrontal cortex tissues were carefully removed on ice. The tissues were weighed quickly, and the total protein extracts of the prefrontal cortex were obtained by homogenization in mammalian cell lysis reagent (Pioneer Biotechnology, Shanghai, China) with a protease inhibitor mixture (Roche, Indianapolis, IN, USA). The levels of IL-6, prostaglandin-endoperoxide synthase 2 (PTGS2), and tumor necrosis factor alpha (TNF-α) were quantified using a QuantiCyto ELISA kit (NeoBioscience, Shenzhen, Guangdong, China). Real-time PCR was performed on tissues from the prefrontal cortexes of mice (see methods above).

### Immunoprecipitation coupled to mass spectrometry (IP-MS) and data analysis

U937 cells were infected by FAM76B-Strep tagⅡ or control Strep tagⅡ-expressing lentiviruses, followed by stable cell line screening. The total protein was extracted, purified with Strep-Tactin beads (QIAGEN, Düsseldorf, Germany), and sent to Shanghai Bioprofile Technology for mass spectrometry sequencing. After high-performance liquid chromatography and mass spectrometry (LC-MS/MS) analysis, the MS data were analyzed using MaxQuant software version 1.6.0.16. MS data were searched against the UniProtKB Rattus norvegicus database (36,080 total entries, downloaded on 08/14/2018). Trypsin was selected as the digestion enzyme. A maximum of two missed cleavage sites and mass tolerances of 4.5 ppm for precursor ions and 20 ppm for fragment ions were defined for the database search. Carbamidomethylation of cysteines was defined as a fixed modification, while acetylation of the protein N-terminal and oxidation of methionine were set as variable modifications for database searching. The database search results were filtered and exported with a <1% false discovery rate at the peptide-spectrum-matched level and the protein level.

### Co-immunoprecipitation

Human cDNAs of full-length hnRNPA2B1 and different domains of hnRNPA2B1, including RRM1, RRM2, and RGD, were constructed by PCR using pGEMT/hnRNPA2B1 (kept in the Xia lab) as the template and the primers listed in Table S1. HEK293 cells were co-transfected with FAM76B-Streptag Ⅱ and hnRNPA2B1-Flag-expressing vectors. At 48 h after transfection, approximately 500 µg of protein extracts prepared from these cells were incubated with Strep-Tactin beads at 4°C for 3 h. The bound protein was examined by Western blot with anti-Flag or anti-FAM76B antibodies. Co-immunoprecipitation of hnRNPA2B1 and IκB-flag was performed similarly, except that protein A/G-Sepharose beads (Thermo, Rockford, IL, USA) charged with anti-hnRNPA2B1 antibody or mouse normal serum were used.

### Confocal microscopy

HEK293 cells expressing the fusion proteins eGFP-FAM76B and mCherry-tagged hnRNPA2B1 were visualized using a Leica TCS-SP8 confocal microscope (Leica Microsystems Inc., Shanghai, China).

### Human tissues

Autopsy brain tissues from patients who had had either a TBI or dementia were collected under an IRB-approved protocol at the University of Utah. Paraformaldehyde-fixed, paraffin-embedded human brain samples from 23 dementia cases (Table S4) were acquired from the Neuropathology Core of Northwestern University’s Center for Cognitive Neurology and Alzheimer’s disease. Demographic and neuropathologic data for these cases are presented in Table S4. Pathologic characterization was made by board-certified neuropathologists blinded to case identity and following consensus criteria (Mackenzie et al. 2009; Cairns et al. 2007; Mackenzie et al. 2010; McKhann et al. 2001).

### Immunofluorescence and immunohistochemistry

For immunofluorescence, unstained slides were deparaffinized, followed by antigen retrieval using Diva Decloaker buffer (Biocare Medical, Pacheco, CA, USA). The slides were then blocked with 1% bovine serum albumin and 0.1% Triton in PBS for 1 hour, before applying primary antibody overnight. After overnight incubation, secondary antibody was applied for 45 minutes, before being washed and coverslipped. The following primary antibodies were used: anti-FAM76B (monoclonal, 1:1000, homemade (Zheng et al. 2016)), IBA-1 (goat polyclonal, 1:1000, Abcam, Boston, MA, USA), or hnRNPA2B1 (rabbit polyclonal, 1:1000, Abcam, Boston, MA, USA) antibodies. Secondary antibodies included donkey anti-rabbit, donkey anti-goat, and donkey anti-mouse antibodies (1:500; Abcam, Boston, MA, USA). Cultured cells were stained in a similar manner except without being deparaffinized and without the antigen retrieval step. For immunohistochemistry, unstained slides were deparaffinized, followed by antigen retrieval using Diva Decloaker buffer. The slides were then quenched with 10% H_2_O_2_ in methanol and blocked as described above. The primary antibodies, including monoclonal anti-FAM76B (monoclonal, 1:1000, homemade (Zheng et al. 2016)) or IBA-1 (goat polyclonal, 1:1000, Abcam, Boston, MA, USA) antibodies, were then applied overnight, followed by biotinylated secondary antibody (1:500; Abcam, Boston, MA, USA). Signal detection was performed using a VECTASTAIN Elite ABC kit and DAB (Vector Laboratories, Burlingame, CA, USA). The stains were reviewed using an Olympus BX53 microscope (Tokyo, Japan). Representative images were taken with an Olympus DP74 camera, and cellSens Dimension software was used for brightness and contrast adjustment and image cropping.

### Microglial quantification

Microglia were counted using a 40× objective with a grid (250 × 250 µm^2^) in a minimum of five microscopic grid fields in the area of interest per slide. Results are given as mean objects per unit area (mm^2^). Microglial cells that had stained cytoplasmic processes and contained a nucleus in the plane of the section were counted.

### Statistics

Group effects were evaluated using unpaired t-tests (Mann-Whitney), paired t-tests, and one-way ANOVA followed by the Tukey honest significant difference (HSD) test. The difference in the ratio of the microglia density was evaluated by Chi-square goodness-of-fit tests. All statistical analyses were performed using GraphPad Prism software (version 4.01). Differences between the means were considered significant at p<0.05.

### Study approval

All animal studies were performed in accordance with institutional guidelines and with approval by the Institutional Animal Care and Use Committee of Shaanxi Normal University (SNNU 2019-0128). Ethical permit of the use of the samples of human brain autopsy specimens was granted by the ethics committee of Shaanxi Normal University.

## 3. Results

### 3.1 FAM76B plays an anti-inflammatory role in macrophages *in vitro*

FAM76B is a nuclear speckle localized protein with previously unknown function. We previously found that FAM76B was highly expressed in U937 cells using a homemade FAM76B antibody (Zheng et al. 2016). U937 is a human macrophage cell line that has been widely used to study inflammation *in vitro*. Therefore, we hypothesized that FAM76B may involve inflammation in U937 cells. To test the hypothesis, we first produced a FAM76B knockdown U937 cell line (*Fam76b* KD) by lentivirus-mediated Cas9/sgRNA genome editing. The expression of FAM76B in U937 cells with *Fam76b* KD was confirmed by Western blot (Fig. 1a). Following 24 h of treatment with PMA plus LPS, the cell line showed markedly increased expressions of IL-6, PTGS2, TNF-α, and IL-10 (Fig. 1b). The increase in IL-6 expression in the *Fam76b* KD cell line was more prominent than the increases in PTGS2 or in TNF-α expression. The results indicated that FAM76B was involved in regulating inflammation in U937 cells. Furthermore, a FAM76B gene knockout U937 cell line (*Fam76b^-/-^)* was obtained using Cas9/sgRNA technology followed by drug screening and dilution cloning, then was confirmed by Western blot and sequencing (Fig. 1c and Fig. S1). Similarly, significantly increased IL-6 expression was observed in the PMA+LPS-treated *Fam76b^-/-^* cell line (Fig. 1d). Moreover, lentivirus-mediated overexpression of FAM76B rescued the function of FAM76B and reduced cytokine mRNA levels in *Fam76b^-/-^* cells (Fig. 1e and 1f). These results demonstrated that FAM76B could inhibit inflammation in macrophages *in vitro*.

**Figure 1.**
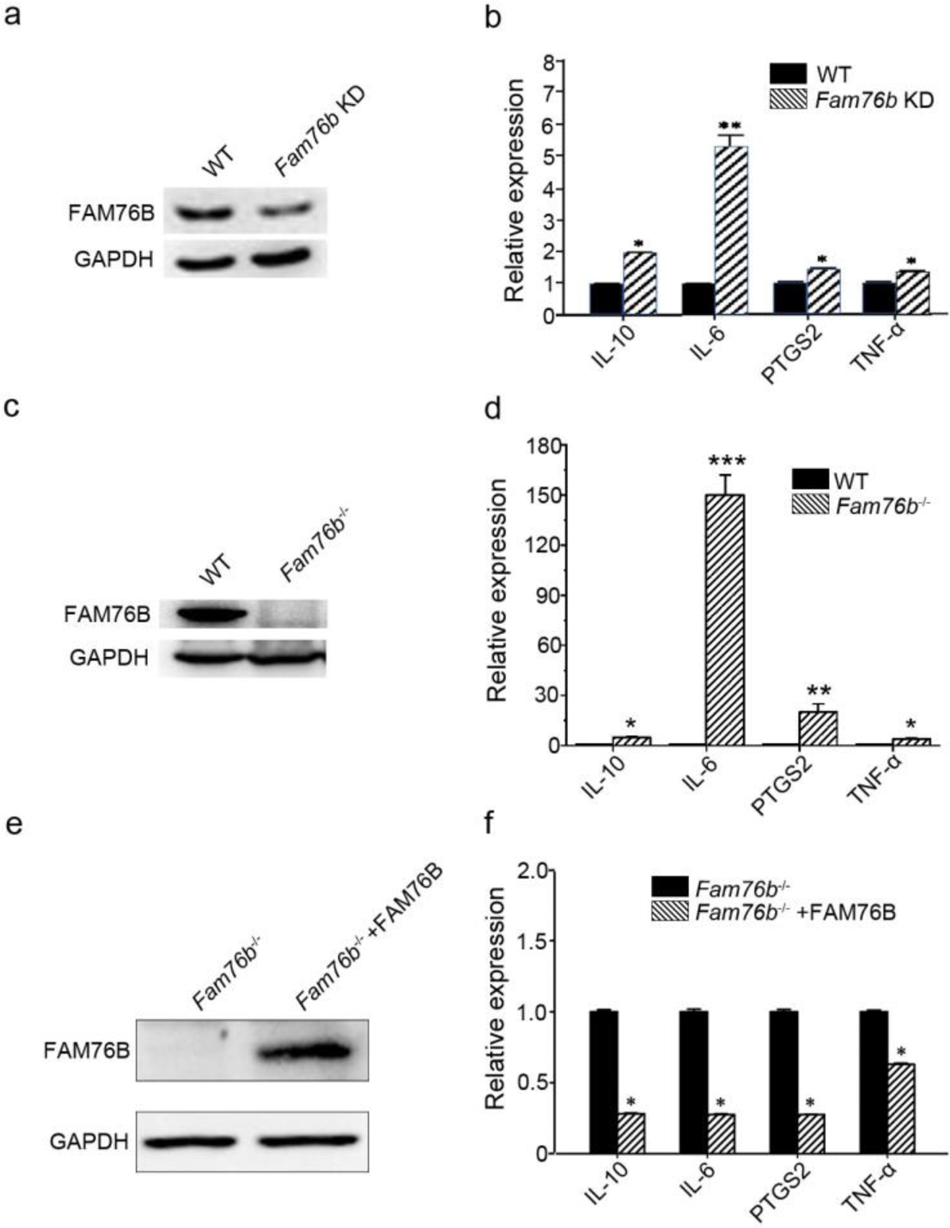
FAM76B regulates the expression of cytokines in U937 cells. (a) Western blot for the detection of FAM76B expression in U937 with FAM76B knockdown cells generated with the Cas9/sgRNA technique. (b) FAM76B knockdown in U937 increased the expression of IL-6, PTGS2, TNF-α, and IL-10, as determined by real-time PCR in the presence of PMA+LPS. (c) Western blot for the detection of FAM76B expression in U937 with FAM76B knockout cell line generated with the Cas9/sgRNA technique. (d) FAM76B knockout in U937 cells significantly enhanced the expression of IL-6, PTGS2, TNF-α, and IL-10, as determined by real-time PCR in the presence of PMA+LPS. (e) Western blot validated the rescued expression of FAM76B in the FAM76B knockout U937 cell line infected with the lentiviral vector expressing FAM76B. (f) The rescued expression of FAM76B in the U937 cell line with FAM76B knockout reduced the mRNA levels of cytokines in the presence of PMA+LPS. The experiments were performed at least three times. *p<0.05, **p<0.01, ***p<0.001, statistically significant.

### 3.2 FAM76B regulates the NF-κB pathway by influencing the translocation hnRNPA2B1 proteins

The data above showed that FAM76B has an anti-inflammatory effect by suppressing the expression of proinflammatory cytokines, especially IL-6. IL-6 is one of the important mediators of the inflammatory response, and it can be activated by NF-κB. NF-κB/IL-6 signaling has long been considered a major proinflammatory signaling pathway in the peripheral tissues and brain. To explore the mechanisms of FAM76B regulating inflammation in U937 cells, the luciferase reporter vector of the IL-6 promoter was constructed. HEK293 cells transfected with this vector showed upregulated activity of the IL-6 promoter by NF-κB overexpression, confirming the activity of the reporter vector (Fig. 2a). Interestingly, FAM76B knockout in HEK293 cells further increased the promoter activity of IL-6, compared to WT cells (Fig. 2a). In WT U937 cells, the activity of the IL-6 promoter was increased after LPS treatment, and FAM76B knockout in U937 cells made this change even more prominent (Fig. 2b). Next, to evaluate if FAM76B regulates IL-6 promoter activity by influencing NF-κB, we constructed luciferase reporters controlled by a miniCMV promoter containing the NF-κB binding motif 1 or 2 sequence from the IL-6 promoter region. The results indicated that the luciferase activity from the vector in both WT and FAM76B knockout U937 cells was increased, especially in the latter, after treatment with 1 ng/ml LPS (Fig. 2c and 2d). These data suggested that FAM76B inhibited the activity of the IL-6 promoter by affecting the NF-κB pathway.

**Figure 2.**
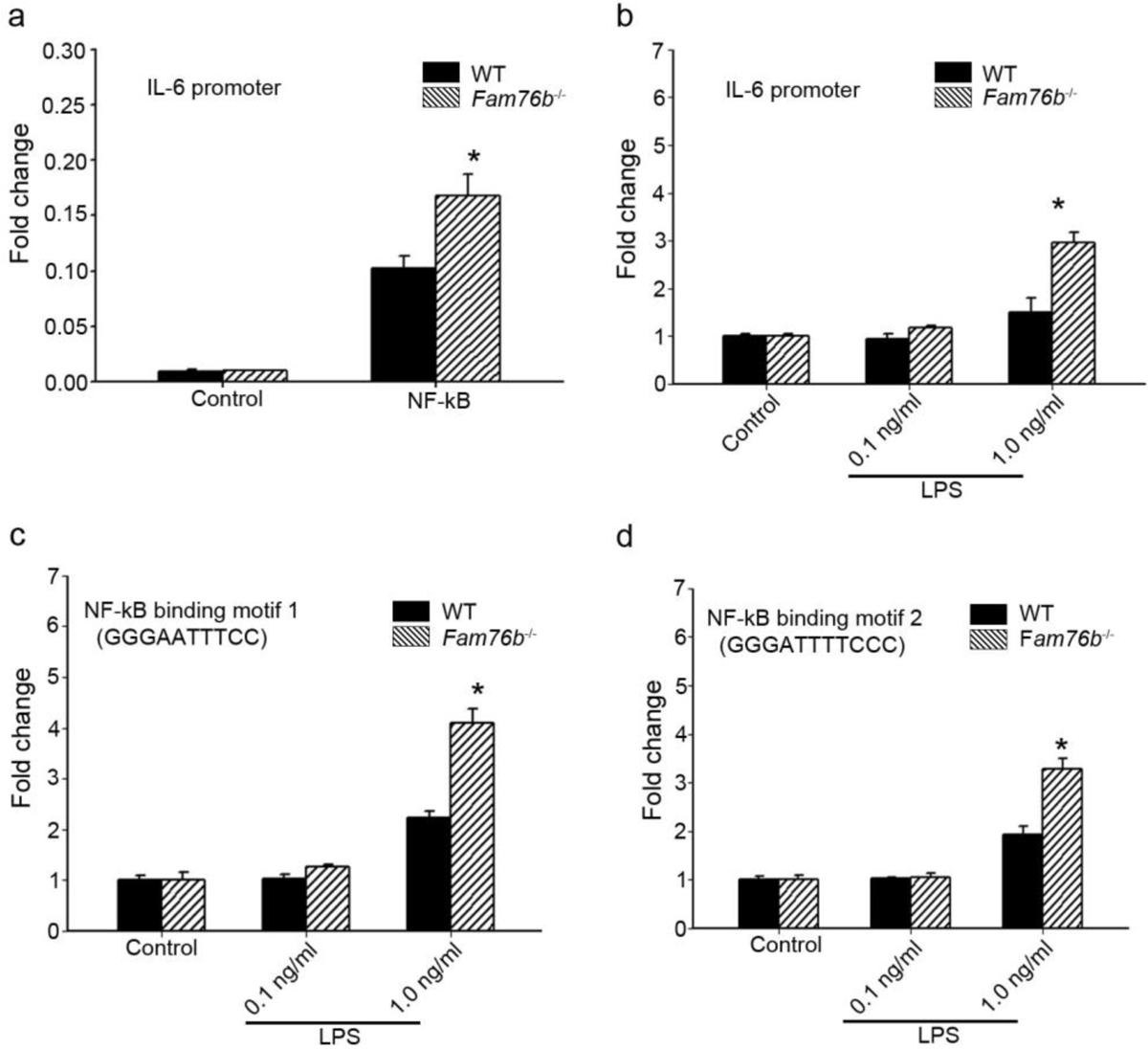
FAM76B regulates IL-6 promoter activity by affecting the NF-κB pathway. (a) The luciferase reporter vector of the IL-6 promoter was tested in WT and FAM76B knockout (*Fam76b*^-/-^) HEK293 cells. The luciferase activity was low in both WT and FAM76B knockout HEK293 cells and significantly increased after transfecting the NF-κB-expressing vector. The increased activity of the IL-6 promoter was more prominent in FAM76B knockout HEK293 cells than in WT cells. (b) The IL-6 promoter was increased in WT and FAM76B knockout U937 cells, but was more prominent in the latter, after LPS treatment. (c and d) Luciferase activity was increased in WT and FAM76B knockout U937 cells carrying NF-κB binding motifs 1 or 2 (and more prominently in the FAM76B knockout U937 cells), indicating that FAM76B inhibited NF-κB binding activity of IL-6 promoter. The experiments were repeated at least three times. *p<0.05, statistically significant.

To further explore the mechanisms of FAM76B regulating inflammation via NF-κB, the FAM76B-interacting proteins were investigated using immunoprecipitation coupled to mass spectrometry (IP-MS) in U937 cells. We identified 160 proteins that interact with FAM76B within U937 cells. Selective molecules of these proteins are listed in Table S5, with their interaction scores and ranks. To validate the IP-MS results, we performed co-immunoprecipitation on selected proteins. Among those FAM76B interacting proteins, hnRNPs captured our attention because of their reported function in regulating the NF-κB pathway (Zhao et al. 2009; Ma et al. 2022). The hnRNPA2B1 was selected for further validation. The co-immunoprecipitation results confirmed the interaction of FAM76B with hnRNPA2B1 (Fig. 3a). In addition, confocal microscopy also revealed the co-localization of FAM76B and hnRNPA2B1 in the nucleus of HEK293 cells overexpressing FAM76B-eGFP and hnRNPA2B1-mCherry (Fig. 3b). Based on the validation of the interaction between hnRNPA2B1and FAM76B, to identify the domain(s) of the hnRNPs required for their interaction with FAM76B, we generated different truncates containing RRM1, RRM2, or RGD domains of hnRNPs and demonstrated that the RGD domain of hnRNPs was responsible for its binding to FAM76B (Fig. 3c and 3d). It has been reported that hnRNPA1 binds to IΚBα, which leads to IκBα degradation and consequently, to NF-κB activation (Zhao et al. 2009). Using co-immunoprecipitation with HEK293 cells transfected with plasmids expressing hnRNPA2B1 and IκBα-flag (or IκBε-flag), we also demonstrated that hnRNPA2B1 binds to IκBα or IκBε (Fig. 3e). Based the results above, we speculated that FAM76B, hnRNP A/B, and IκBs could form a protein complex by binding FAM76B to hnRNPs through the RGD domain and binding hnRNPs to IκBs through the RRM2 domain (Fig. 3f). The results above suggested that FAM76B could regulate inflammation by its interaction with hnRNPA2B1, which then leads to IκBα degradation by its interaction with IκBα, followed by NF-κB activation.

**Figure 3.**
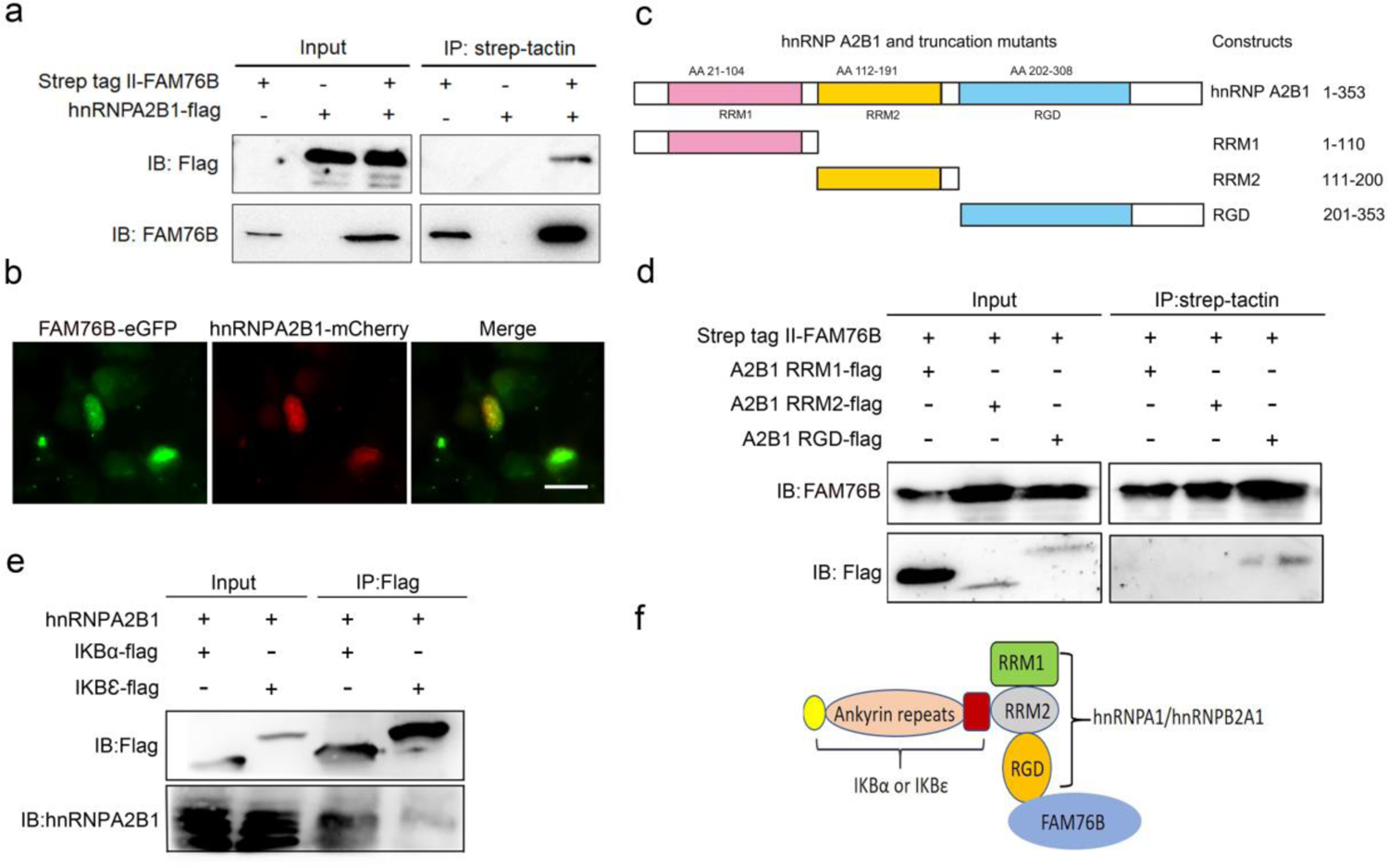
Validation of the interactions among FAM76B, hnRNPA2B1, and IκBs. (a) The interaction between FAM76B and hnRNPA2B1 was revealed by co-immunoprecipitation. FAM76B-Strep-tag Ⅱ and hnRNPA2B1-Flag were overexpressed in HEK293 cells, followed by co-immunoprecipitation of FAM76B and hnRNPA2B1 using whole cell lysate and Western blot with anti-Flag or anti-FAM76B antibodies. (b) Confocal microcopy revealed the co-localization of FAM76B and hnRNPA2B1 in the nucleus of HEK293 cells transfected with plasmids expressing FAM76B-eGFP and hnRNPA2B1-mCherry, respectively. Scale bar, 20 µm. (c) An illustration of hnRNPA2B1 domains (RRM1, RRM2, and RGD) tagged with Flag generated for detecting the hnRNPA2B1 region(s) responsible for binding FAM76B. (d) Identification of the hnRNPA2B1 domains responsible for binding to FAM76B. Strep tag Ⅱ-FAM76B and hnRNPA2/B1 domain-flag were transfected into HEK293 cells, followed by co-immunoprecipitation and Western blot with anti-Flag or anti-FAM76B antibodies, which showed the interaction between the RGD domain of hnRNPA2B1 and FAM76B. (e) Similarly, the interaction between hnRNPA2B1 and IKBα-flag or IκBε-flag was detected by co-immunoprecipitation. (f) Schematic diagram of FAM76B, hnRNPA2B1, and IKBs’ protein complex formation: hnRNPA2B1 binds FAM76B by its RGD domain and binds IκBs by its RRM2 domain.

FAM76B and hnRNPA2/B1 are nuclear localized proteins; however, it has been reported that hnRNPA2/B1 could translocate from the nucleus to the cytoplasm, which would lead to activation of the NF-κB pathway (Wang et al. 2019). Considering the interaction of FAM76B and hnRNPA2B1, we speculated that FAM76B is the protein that mediates the cytoplasmic translocation of hnRNPA2B1, which is then followed by NK-κB pathway activation. To test that hypothesis, hnRNPA2B1 immunostaining was performed on U937 cells with *FAM76B* knockout, which showed increased cytoplasmic translocation of hnRNPA2B1 (Fig. 4a); this result was also confirmed by the levels of nuclear and cytoplasmic hnRNPA2B1 using Western blot and semiquantification based on the results of Western blot (Fig. 4b and 4c). Furthermore, we found that *FAM76B* knockout in U937 cells resulted in increased phosphorylation of endogenous IKKα, IKKβ, and the downstream molecule IκBα upon LPS stimulus (Fig. 4d), which was also confirmed by semiquantification (Fig. 4e). In addition, we observed a concurrent increase in the nuclear translocation of p65, a hallmark of classical NF-κB pathway activation, by Western blot and semiquantification in LPS-stimulated U937 cells with *Fam76b* knockout (Fig. 4f and 4g). Interestingly, when U937 was induced into the M1-like macrophage state with PMA followed by LPS+IFNγ stimulation, both FAM76B and hnRNPA2B1 were found to be partially translocated into the cytoplasm from the nucleus (Fig. 4h). Together, these results indicated that FAM76B could promote NF-κB activation by affecting the translocation of hnRNPA2B1.

**Figure 4.**
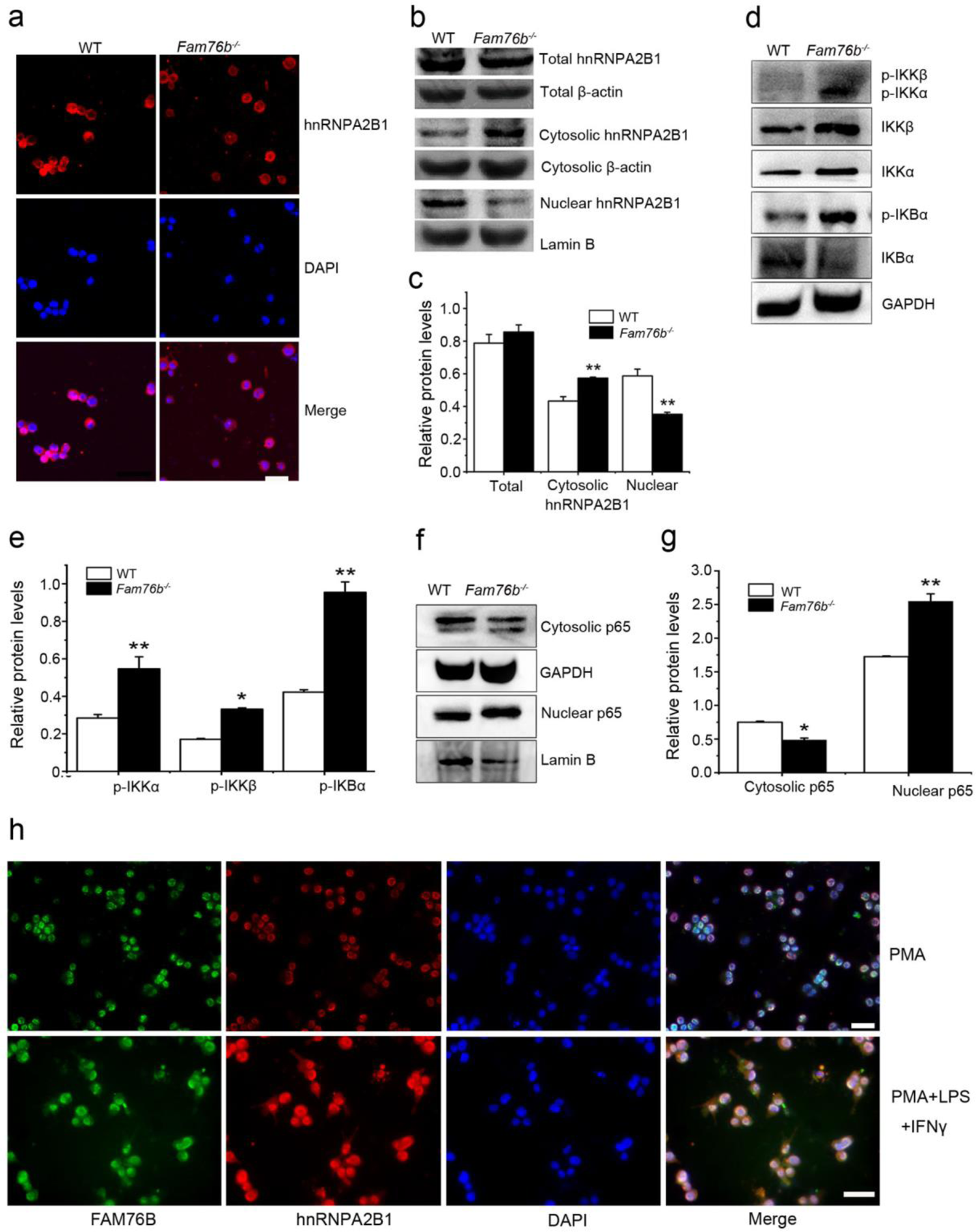
FAM76B regulates the NF-κB pathway by influencing the translocation of hnRNPA2B1. (a) Immunofluorescence revealed increased cytoplasmic translocation of hnRNPA2B1 in U937 cells with *FAM76B* knockout (*FAM76B^-/-^* U937) stimulated with PMA for 48 h. Scale bar, 20 µm. (b) Western blot confirmed the cytoplasmic translocation of hnRNPA2B1 in U937 cells with *FAM76B* knockout (*FAM76B^-/-^* U937) stimulated with PMA for 48 h. (c) The semiquantification of the results of Western blot from Fig. 4b. (d–e) Western blot revealed the increased phosphorylation of endogenous IKKα, IKKβ, and IκBα in *FAM76B^-/-^* U937 cells stimulated with LPS. (f) The semiquantification of the Western blots result from Fig. 4d and 4e. (g) Western blot showed the increased nuclear translocation of p65 in *FAM76B^-/-^* U937 cells stimulated with PMA followed by incubation with LPS and hIFNγ. (h) The semiquantification of the Western blot result from Fig. 4g. The experiments were performed at least three times. *p<0.05, **p<0.01, statistically significant. (i) Stimulation with 1 ng/ml PMA followed by LPS and IFNγ treatment leads to the cytoplasmic translocations of FAM76B and hnRNPA2B1 in U937 cells. Scale bar, 20 µm.

### 3.3 Inflammation mediated by macrophages and microglia are enhanced in *Fam76b* knockout C57BL/6 mice

Our previous study showed that FAM76B is widely expressed in different human organs, with the highest expression levels found in the brain and spleen (Zheng et al. 2016). Similarly, here we found by real-time PCR that the mouse brain and spleen also had high levels of FAM76B expression (Fig. S2), which suggests that FAM76B may have important functions related to inflammation in these tissues. To investigate whether FAM76B has anti-inflammatory activity *in vivo,* we produced *Fam76b* knockout C57BL/6 mice (*Fam76b^-/-^*) by gene-trap technology (Fig. 5a). FAM76B knockout was confirmed by genotyping analysis and the expression of FAM76B at the mRNA and protein level (Fig. 5b–d). *Fam76b^-/-^* mice showed normal weight gain and lifespan (Fig. S3). *Fam76b^-/-^* mice developed spleens that were visibly enlarged and weighed more than those of WT mice (Fig. 6a and 6b). Hematoxylin and eosin (H&E)-stained sections of the spleen showed normal red pulp and hypertrophy of white pulp (Fig. 6a). Flow cytometry revealed an increase in the CD11b+ myeloid population and a slight decrease in CD19+ B cells in *Fam76b^-/-^* mice, while no changes were found in CD3+ T cells (Fig. 6c–g). To further elucidate the effect of FAM76B on the function of macrophages in the *Fam76b^-/-^* mice, we examined their spleens following intraperitoneal LPS injection. There were no significant morphologic changes in the WT and knockout mice (Fig. 6h). However, after LPS treatment, the knockout mice had many tingible body macrophages in the white pulp of the spleens, while the WT mice had none (Fig. 6h). These data suggested that loss of FAM76B expression might lead to a decreased ability of the white pulp macrophages to clear out the apoptotic cells produced during the germinal center reaction. The results above indicated that FAM76B could be involved in macrophage function.

**Figure 5.**
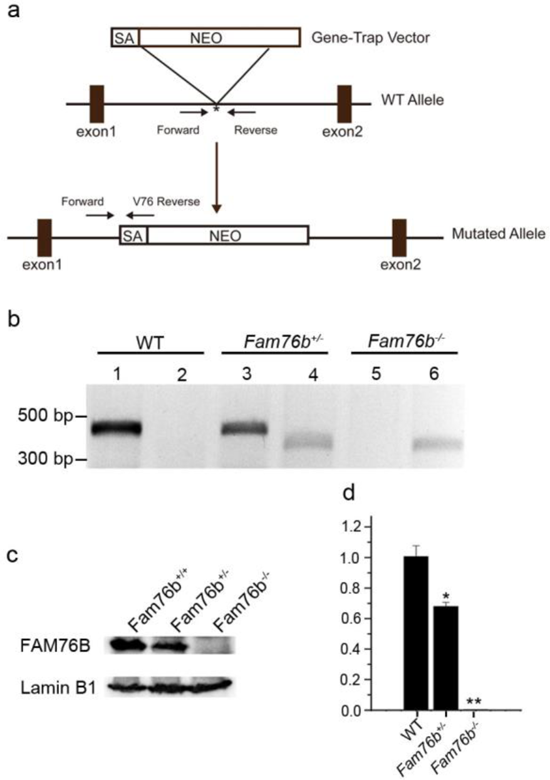
Generation of *Fam76b* knockout (*Fam76b^-/-^*) mice. (a) Schematic diagram of the homologous recombination construct for generating *Fam76b^-/-^* mice, with arrows denoting primer locations. (b) Genotyping results for Fam76b WT, hemizygous (+/-), and homozygous (-/-) mice. Lanes 1, 3, and 5 are the amplified products of Fam76b forward and reverse primers (Table S2); lanes 2, 4, and 6 are the amplified products of Fam76b forward and V76 reverse primers (Table S2). (c) Western blot confirmed FAM76B protein levels in mouse embryonic fibroblasts from Fam76b WT, hemizygous (+/-), and homozygous (-/-) mice. (d) Real-time PCR of Fam76b mRNA levels in mouse embryonic fibroblasts from Fam76b WT, hemizygous (+/-), and homozygous (-/-) mice. *p<0.05, **p<0.01, statistically significant.

**Figure 6.**
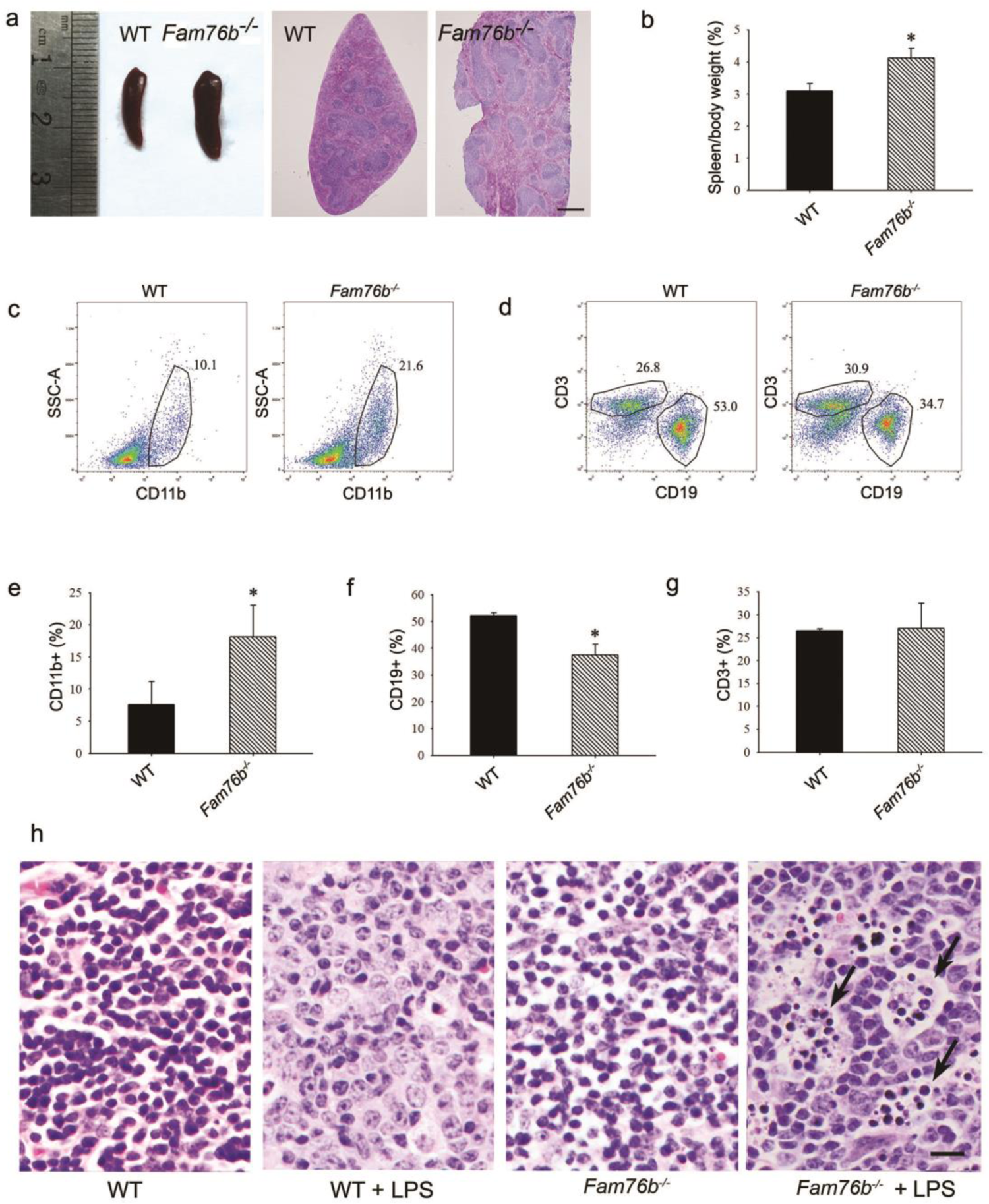
*Fam76b* knockout mice had enlarged spleens with altered cell populations and inflammation. (a) Enlarged spleen with white pulp hypertrophy in *Fam76b* knockout mice (*Fam76b*^-/-^) (5 months), as compared with WT. Scale bar, 500 µm. (b) The weight ratio of spleen to body in *Fam76b* knockout mice (*Fam76b*^-/-^) (5 months) revealed an enlarged spleen, as compared with WT. *p<0.05, statistically significant. (c–g) Flow cytometry results of the cell population of spleens of *Fam76b*^-/-^ mice (5 months). Dot plots (c and d) and bar graphs (e, f, and g) show increased populations of CD11b+ and decreased populations of CD19+ B cells in *Fam76b*^-/-^ spleens (n=4; 5 months) (*p<0.05, one-way ANOVA). (h) Photomicrographs of H&E-stained spleens from *Fam76b*^-/-^ mice intraperitoneally injected with LPS show abundant tingible body macrophages (arrows) in the germinal center, whereas no tingible body macrophages were seen in LPS-treated WT or in phosphate-buffered saline (PBS)-treated *Fam76b*^-/-^ and WT mice. Scale bar, 25 µm.

Macrophages play an important role in inflammation, so to further investigate the effect of FAM76b on the regulation of macrophage-mediated inflammation *in vivo*, BMDMs from *Fam76b^-/-^* mice were isolated and used to analyze IL-6 expression. The results indicated that macrophages from FAM76B knockout mice could significantly increase IL-6 expression compared to WT murine macrophages (Fig. S4a). Moreover, the increased expression of IL-6 could be downregulated when FAM76B was rescued in macrophages from FAM76B knockout mice (Fig. S4b), which was consistent with the results obtained from FAM76B knockout U937 cells *in vitro*. The results above indicated that FAM76B possesses anti-inflammation activity *in vivo*.

Microglia play a crucial role in mediating neuroinflammation. Therefore, we tested whether deleting FAM76B could enhance inflammation mediated by microglia in FAM76B knockout mice. First, immunostaining with anti-IBA-1 antibody (Ito et al. 1998; Okere et al. 2000; Hirayama et al. 2001) was used to reveal the total microglial population in the hippocampus and thalamus of Fam76b^-/-^ mice.IBA1-positivity microglia were higher in 12-month-old Fam76b knockout mice than that in age-matched WT mice, and displayed a highly reactive morphologies with abundant cytoplasm (Fig.7a). However, there are no significant difference between 4-month-old Fam76b-/- and WT mice (data not shown). An evaluation of the densities of IBA-1-positive microglia (number/mm^2^) showed more IBA-1-positive microglial cells in the CA1 region and thalamus of 12-month-old Fam76b^-/-^ mice than in those same brain areas of age-matched WT mice. The densities of IBA-1-positive microglia were similar between 4-month-old Fam76b^-/-^ and WT mice (Fig. 7b). A CCI mouse model (Zheng et al. 2022) was then used to examine the effect of FAM76B on microglia-mediated neuroinflammation. The densities of IBA-1-positive microglia were higher in the hippocampus adjacent to the cortical contusion of Fam76b^-/-^ mice than in that of WT mice or sham controls (Fig. 7c and 7d). Real-time PCR showed that the expression level of IL-6 in the ipsilateral hippocampus was elevated in both Fam76b^-/-^ and WT mice 3 days after TBI, with more prominent changes observed in Fam76b^-/-^ mice (Fig. 7e). We then stereotactically injected LPS into the left frontal lobes of Fam76b^-/-^ and WT mice. Seven days post-treatment, histologic examination revealed abundant macrophages at the injection site of Fam76b^-/-^ mice, but only a mild inflammatory response in WT mice (Fig. S5a). Real-time PCR and ELISA showed that the expression level of IL-6 was elevated in both WT and Fam76b^-/-^ mice 24 h post-LPS treatment, with more prominent changes evident in the knockout mice (Fig. S3b and S3d). TNF-α expression was similarly altered as in the WT and Fam76b^-/-^ mice, but to a lesser extent than IL-6 (Fig. S5c and S5e). These experiments indicate that FAM76B is also involved in modulating neuroinflammation mediated by microglial cells *in vivo*.

**Figure 7.**
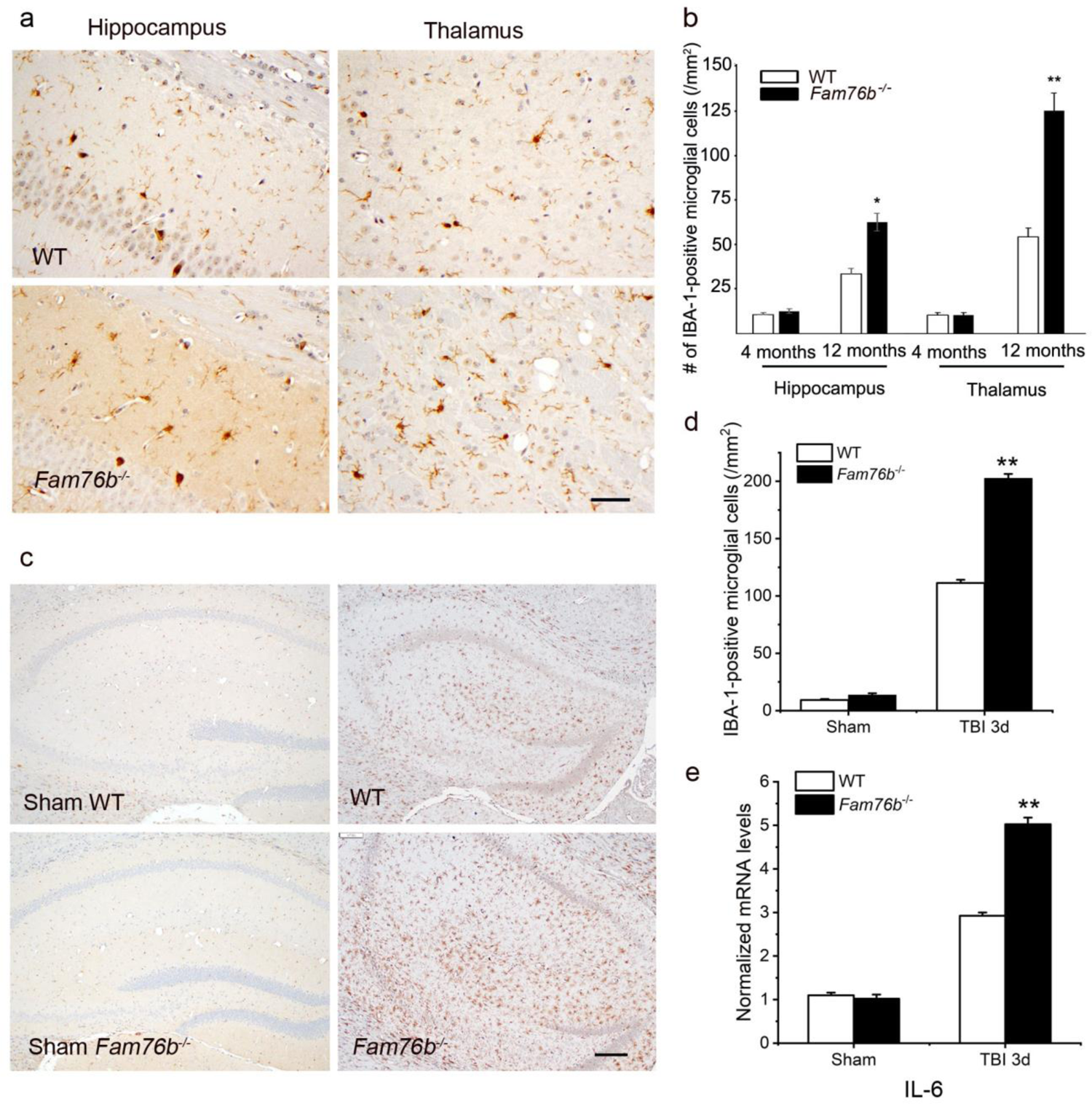
Neuroinflammation was enhanced in *Fam76b* knockout mice. (a) Increased IBA-1-positive microglial infiltration in the hippocampus (lower left panel) and thalamus (lower right panel) of 12-month-old *Fam76b* knockout mice (*Fam76b^-/-^)*, as compared with age-matched WT mice. Scale bar, 50 µm. (b) Density of IBA-1-positive microglia in hippocampal CA1 regions and the thalamus of 4- and 12-month-old *Fam76b^-/-^* mice, as compared with age-matched WT mice. (c) Increased IBA-1-positive microglia in the hippocampus adjacent to the contusion site of *Fam76b* knockout mice (*Fam76b^-/-^)*, as compared with WT mice and sham controls. Scale bar, 200 µm. (d) Density of IBA-1-positive microglia in hippocampal CA1 regions of *Fam76b^-/-^* mice, as compared with WT mice and sham controls. Values are mean ± SD. The densities of IBA-1-positive microglia were compared to the control value by Student’s t-test (**p<0.01). (e) Increased IL-6 expression, as revealed by real-time PCR, in the ipsilateral hippocampus in both *Fam76b^-/-^* and WT mice 3 days after TBI, with more prominent changes in *Fam76b^-/-^* mice. *p<0.05, **p<0.01, statistically significant.

### 3.4 FAM76B plays a role in neuroinflammation in the context of TBI and neurodegeneration

The results above indicated that FAM76B regulated neuroinflammation in a mouse model *in vivo*. Neuroinflammation is known to be closely related the development of TBI and neurodegeneration. Our previous study showed that FAM76B is widely expressed in different human organs, with the highest expression levels found in the brain and spleen (Zheng et al. 2016). To assess whether FAM76B is involved in the neuroinflammation associated with the human diseases named above, we compared the distribution and expression of FAM76B in diseased brain tissues to those of normal brains. In the normal brain, immunohistochemical stains revealed weak cytoplasmic staining of FAM76B in neurons and nuclear staining in glial cells (Fig. S6a and 6b). In areas of organizing necrosis in brains with TBI, we found that macrophages—which were labeled by the microglial/macrophage marker IBA-1 (Ito et al. 1998; Okere et al. 2000; Hirayama et al. 2001)—were all strongly immunopositive for FAM76B (Fig. S6c). In the hippocampal CA1 area of brains with acute ischemic injury, FAM76B immunostains highlighted many reactive microglial cells (Fig. S6d). These findings strongly suggest that microglial FAM76B is upregulated in injured brains and thus may have important functions in regulating neuroinflammation.

TBI elicits neuroinflammation, which is essential for proper tissue regeneration and recovery (Simon et al. 2017). Thus, we next studied the role of FAM76B in TBI-induced neuroinflammation by examining the expression and cytological distribution of FAM76B/hnRNPA2B1 in human brains with TBI. In normal brain tissue, FAM76B is mainly localized in the nuclei of glial cells, including microglial (Fig. 8a) and oligodendroglial (Fig. S5) cells. hnRNPA2B1 was found to be co-localized with FAM76B in the nuclei of these cells (Fig. 8b). In acute and organizing TBI, in response to the contusion, there was increased expression and cytoplasmic translocation of microglial FAM76B (Fig. 8a), which was co-localized with hnRNPA2B1 (Fig. 8b). In chronic TBI, the residual macrophages/microglia in the previously injured area showed a persistent cytoplasmic FAM76B/hnRNPA2B1 distribution; moreover, the FAM76B- and hnRNPA2B1-positive microglia appeared dystrophic in morphology (Fig. 8a and 8b). These results were consistent with those obtained in U937 cells, supporting the conclusion that FAM76B affects hnRNPA2B1’s translocation from the nucleus to the cytoplasm. These results also demonstrated the activation, evolution, and persistence of FAM76B-positive microglia and the significant role of FAM76B-NF-κB in the organizing process of human TBI.

**Figure 8.**
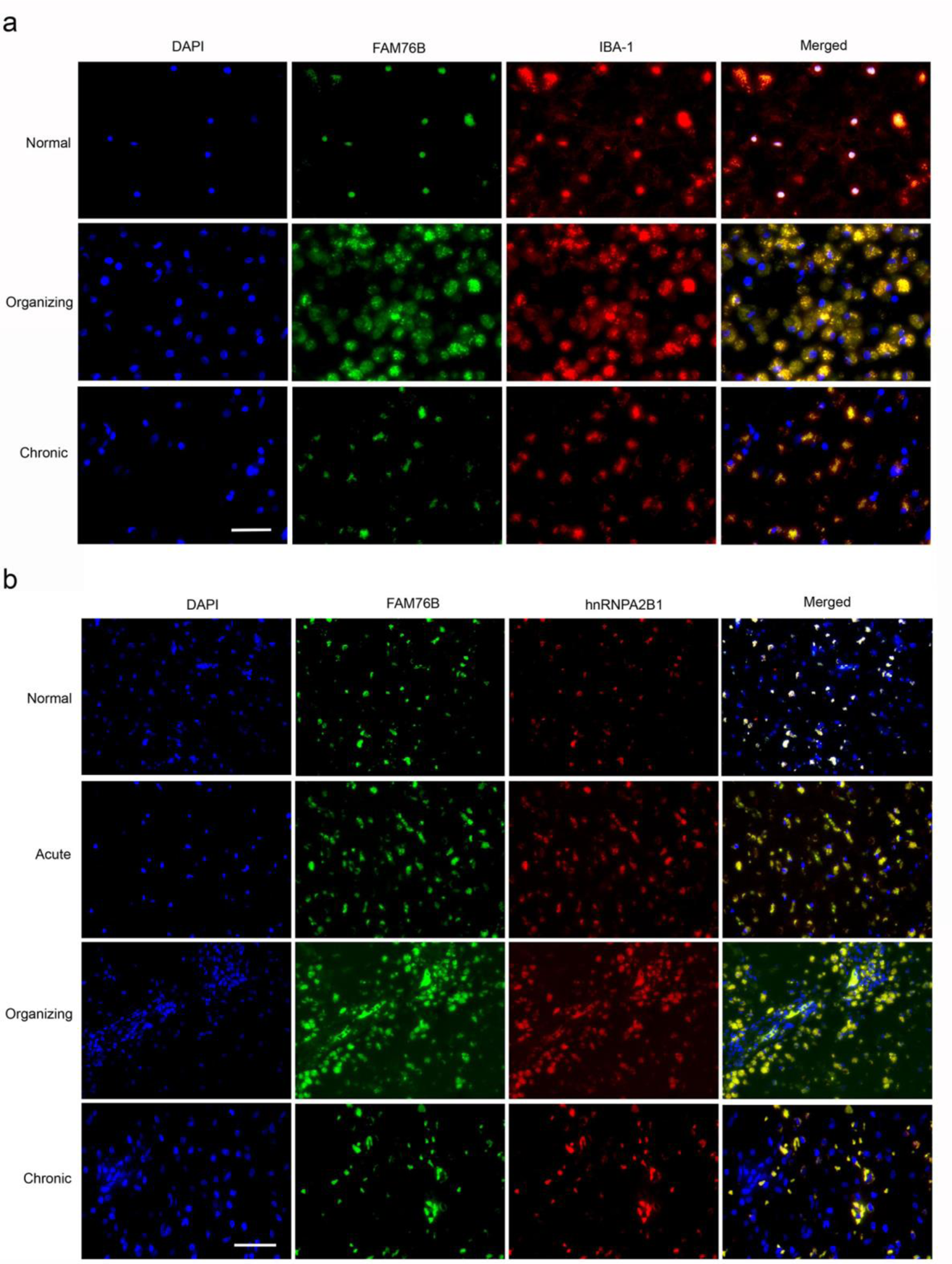
The expression and distribution of FAM76B/hnRNPA2B1 in human brains with TBI. (a)Microglial localization of FAM76B in organizing and chronic TBI. Immunofluorescence revealed the nuclear localization of FAM76B in IBA-1-positive microglia in the normal human cortex. FAM76B was upregulated and cytoplasmically translocated in microglia/macrophages in the human cortex with organizing TBI. In chronic TBI, microglial FAM76B showed persistent cytoplasmic distribution. Scale bar, 100 µm. (b) Upregulation and cytoplasmic translocation of FAM76B and hnRNPA2B1 in acute, organizing, and chronic TBI. Immunofluorescence revealed the nuclear co-localization of FAM76B and hnRNPA2B1 in the normal human cortex. Both FAM76B and hnRNPA2B1 were upregulated and cytoplasmically translocated in microglia/macrophages in the cortex of a patient with acute TBI. Both proteins were further upregulated in the cytoplasm of the microglia/macrophages in the human cortex with organizing TBI. In chronic TBI, microglial FAM76B and hnRNPA2B1 showed persistent cytoplasmic distribution and co-localization. Scale bar, 100 µm.

Neuroinflammation is an important contributor to neurodegeneration. Hence, we evaluated the role of FAM76B in the common neurodegenerative diseases Alzheimer’s disease (AD), frontotemporal lobar degeneration with tau pathology (FTLD-tau), and frontotemporal lobar degeneration with TAR DNA-binding protein 43 inclusions (FTLD-TDP). Similar to what we observed in chronic TBI, FAM76B and hnRNPA2B1 were co-localized in the microglial cytoplasm of brain tissue from patients with neurodegeneration, and the FAM76B- and hnRNPA2B1-positive microglia appeared dystrophic in morphology (Fig. 9a). In addition, immunostains revealed that IBA-1- and FAM76B-positive microglia were scattered in the frontal cortex of normal aging controls and were slightly more numerous in that of AD, FTLD-tau, and FTLD-TDP patients (Fig. 9b). Moreover, the cellular densities of FAM76B-positive and IBA-1-positive microglia were both higher in AD, FTLD-tau, and FTLD-TDP patients than in normal aging controls. (IBA-1-positive microglia, **p<0.01; FAM76B-positive microglia, ^##^p<0.01) (Fig. 9c). The ratio of FAM76B-positive microglial density to IBA-1-positive total microglia density was the highest in the cortex in FTLD-TDP patients (^&&^p<0.01) (Fig. 9c). These data suggest that the FAM76B-NF-κB pathway is activated in AD, FTLD-tau, and FTLD-TDP brains, but is most prominent in FTLD-TDP.

**Figure 9.**
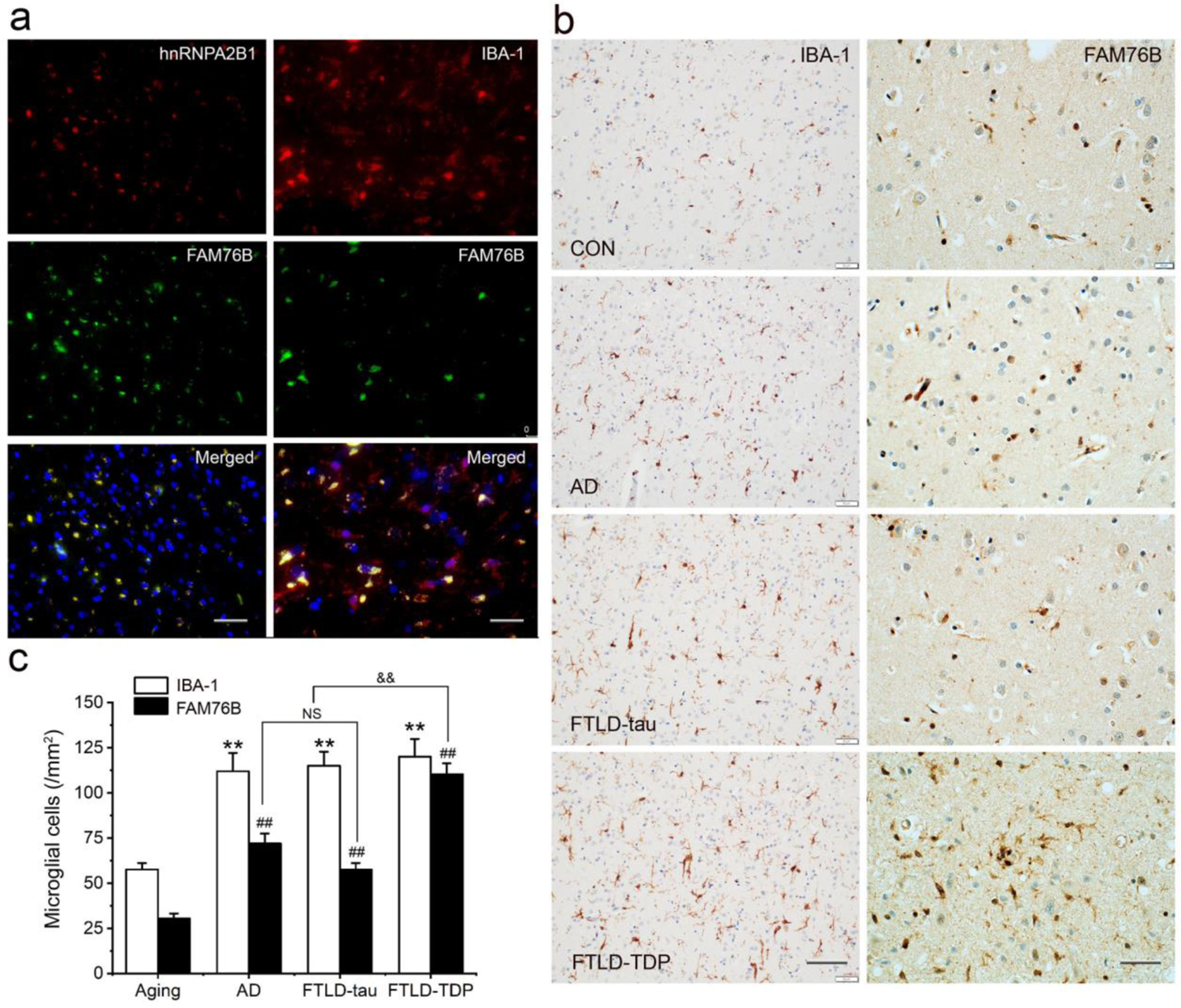
Persistent microglial FAM76B expression in neurodegenerative diseases. (a) Immunofluorescent staining using the frontal cortex of a FTLD-TDP patient demonstrated that FAM76B co-localizes with IBA-1, while FAM76B co-localizes with hnRNPA2B1, in the cytoplasm of microglia. Scale bar, left panels, 100 µm; right panels, 50 µm. (b) Immunohistochemical stains revealed that the frontal cortex of AD, FTLD-tau, and FTLD-TDP patients showed increased IBA-1- and FAM76B-positive microglia as compared to the control (CON). This increase in microglial FAM76B expression was more prominent in FTLD-TDP than in AD or FTLD-tau. Scale bar, left panels for IBA-1 stains, 100 µm; right panels for FAM76B stains, 50 µm. (c) The density of IBA-1- and FAM76B-positive microglia in the frontal cortex of normal aging controls, and AD, FTLD-tau, and FTLD-TDP patients. Values are mean ± SD; n=5. The densities of IBA-1-positive microglia were compared between different groups by one-way ANOVA followed by the Tukey-HSD test (**p<0.01). Similarly, the density of FAM76B-microglia was compared between groups (^##^p<0.01). The ratios of FAM76B- to IBA-1-microglia densities were compared between different groups by Chi-square goodness-of-fit test (^&&^p<0.01). n.s., no significance.

## Discussion

The key findings of this study are as follows. (a) We found that FAM76B is an inflammatory and neuroinflammatory modulator that inhibits NF-κB activity in macrophages/microglia. (b) An interacting partner of FAM76B is hnRNPA2B1, and the cytoplasmic translocation of FAM76B and hnRNPA2B1 was associated with NF-κB pathway activation. (c) BMDMs from FAM76B knockout mice showed an increased inflammatory response in the presence of LPS. FAM76B knockout mice showed dramatically increased tingible body macrophages in the white pulp of the spleen after a local LPS challenge. The brains of FAM76B knockout mice showed age-related neuroinflammation in their hippocampus and thalamus. A local LPS challenge and TBI led to a significantly increased neuroinflammation. (d) Human brains with TBI showed the activation, evolution, and persistence of FAM76B-NF-κB-mediated neuroinflammation during the TBI repair process. (e) There is chronic activation of the FAM76B-NF-κB pathway in dementia, particularly in FTLD-TDP.

This is the first report investigating the function of FAM76B in inflammation and neuroinflammation. We speculate that when FAM76B is present in the nucleus, the hnRNPA2B1 protein is trapped in the nucleus by its binding to FAM76B; however, when FAM76B was decreased or made absent (such as by FAM76B knockdown or knockout, respectively) or was subjected to inflammatory stimulation (such as by LPS), the hnRNPA2B1 protein translocated to the cytoplasm, which then led to increased NF-κB-mediated inflammation by degrading IKBα and causing p65 to enter the nucleus (Fig. 10).

**Figure 10.**
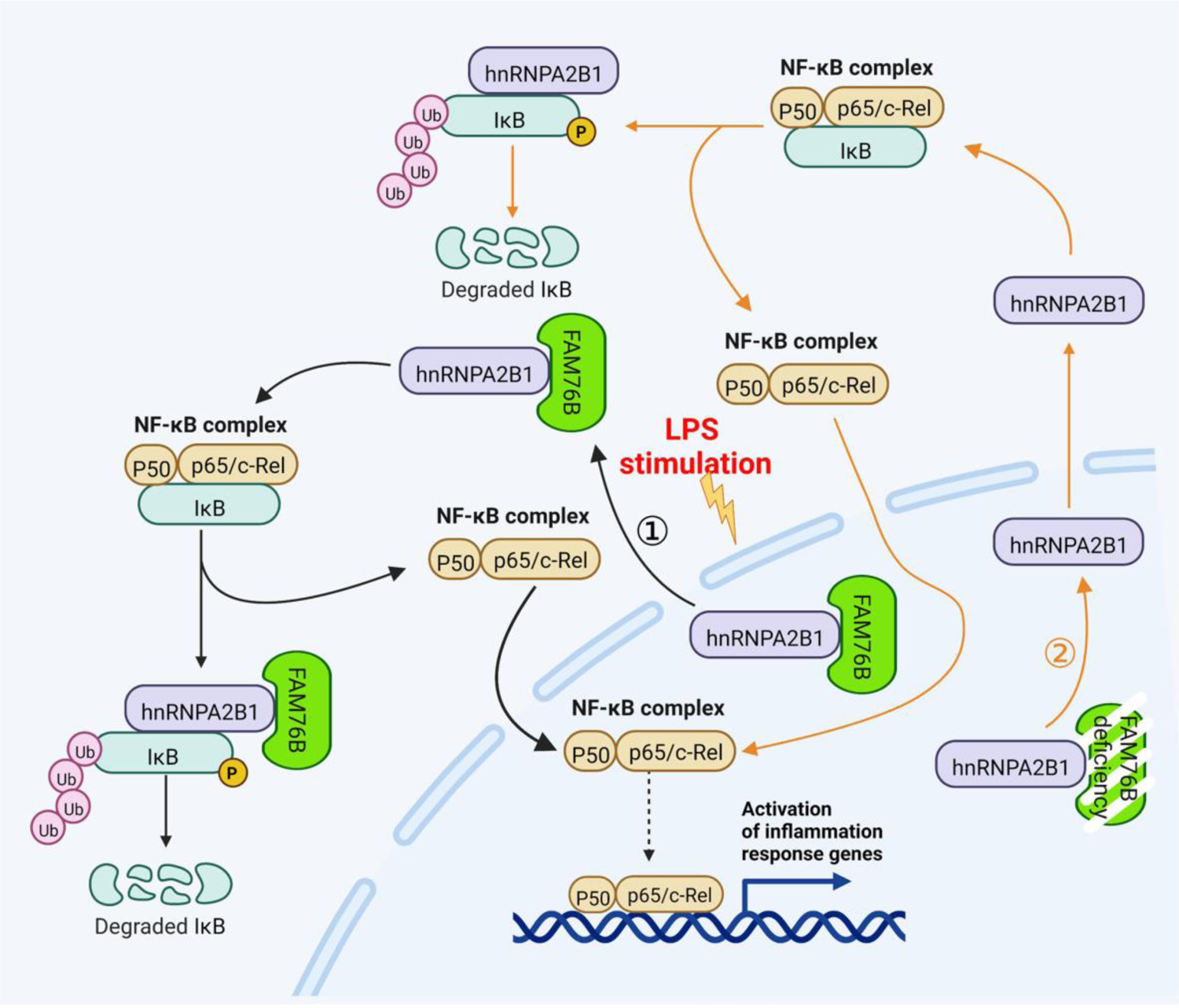
Schematic diagram of FAM76B regulating the NF-κB-mediated inflammatory pathway by affecting the hnRNPA2B1 translocation. Under normal conditions, FAM76B can bind to hnRNPA2B1 and make hnRNPA2B1 stay in the nucleus. However, when FAM76B’s location or expression level of was changed, the localization of hnRNPA2B1 was also changed, which then regulates inflammation in immune cells. ① Upon stimulation with LPS, the FAM76B located in the nucleus of immune cells (such as macrophage or U937 induced macrophage like M1) moved into the cytoplasm, and hnRNPA2B1 was accordingly translocated into the cytoplasm, resulting in enhanced NF-κB-mediated inflammation by the degradation of IKBα and p65’s entry into the nucleus. ② When FAM76B expression was decreased or knocked out in the immune cells, hnRNPA2B1 translocated into the cytoplasm, which led to increased NF-κB mediated inflammation by the degradation of IKBα and p65’s entry into the nucleus.

FAM76B is one of the 86 proteins in the human genome that contains stretches of five or more histidines (Salichs et al. 2009). Studies have suggested that His-repeats may act as nuclear speckle-targeting signals (Alvarez et al. 2003; Herrmann et al. 2001; Salichs et al. 2009). In our previous study, we confirmed the nuclear speckle localization of both human and mouse FAM76B (Zheng et al. 2016). Hence, FAM76B may have functions related to nuclear speckles, such as splicing factor storage and modification (Salichs et al. 2009; McGlincy et al. 2010). We found in this study that FAM76B interacts with hnRNPA2B1, a heterogeneous nuclear ribonucleoprotein related to mRNA binding and splicing (Peng et al. 2021; Moran-Jones et al. 2005) that is associated with inflammation (Lin et al. 2020; Chen et al. 2020; Coppola et al. 2019; Hoffmann et al. 2011). By *in vitro* and *in vivo* experiments, we showed that FAM76B could regulate NF-κB mediated inflammation by influencing the translocation of hnRNPA2B1.

Neuroinflammation is the response of the CNS to injury and disease and is the common thread that connects brain injuries to neurodegenerative diseases (Gilhus et al. 2019; Brambilla. 2019). In TBI, neuroinflammation is one of the most prominent reactions. TBI leads to early resident microglial activation, which is accompanied by local upregulations of TNF-α (Frugier et al. 2010; Csuka et al. 1999) and IL-6 (Frugier et al. 2010; Perez-Barcena et al. 2011; Helmy et al. 2011). Consistently, we also found increased levels of IL-6 at the contusion site of mouse brains after TBI. In addition, the IL-6 level was further increased by FAM76B knockout, indicating the role of the FAM76B-NF-κB pathway in regulating microglial function and driving acute post-traumatic neuroinflammation. TBI can cause persistent neuroinflammation and microglial activation (Simon et al. 2017; Morganti-Kossmann et al. 2019). Studies of TBI biomarkers in adults with severe TBI have shown that serum levels of IL-1β, IL-6, and TNF-α are chronically increased (Simon et al. 2017). TBI animal models have demonstrated persistently increased numbers of microglia at the margins of the lesion and in the thalamus at one year post-injury (Simon et al. 2017). Consistent with these findings, we found that in human brain tissue of chronic TBI patients, there was persistent FAM76B-NF-κB pathway activation in microglia. Persistent microgliosis after TBI correlates with chronic neurodegeneration and dementia development (Simon et al. 2017; Alberici et al. 2018), which suggests that chronic neuroinflammation may be the mechanism of the neurodegeneration associated with TBI. Furthermore, TBI and neurodegeneration may share a similar neuroinflammatory pathway. In this study, we observed that chronic activation of the FAM76B-NF-κB pathway occurs in both chronic TBI and neurodegenerative disorders, particularly FTLD-TDP, suggesting that the FAM76B-NF-κB pathway might be the common pathway that mediates the neuroinflammation in TBI and FTLD-TDP and that FAM76B-mediated neuroinflammation might be the mechanism by which TBI is linked to neurodegeneration.

In the peripheral tissues, FAM76B deficiency led to increased activation of the NF-κB pathway. In response to an acute proinflammatory stimulus, intraperitoneal LPS administration, FAM76B knockout mice showed an interesting phenotype: significantly increased tingible body macrophages in the spleen’s white pulp. This finding suggests that FAM76B plays a unique role in regulating macrophage function. An overlap between FTLD and autoimmune disease has been noted in the field of neurodegeneration (Alberici et al. 2018). FTLD-associated genetic variants are also linked to autoimmune conditions (Bright et al. 2019; Miller et al. 2013; Miller et al. 2016). Hence, FAM76B dysfunction might be involved in autoimmune processes in FTLD. Though the function of FAM76B in the peripheral tissue is beyond the scope of this paper, it is an area that deserves further study.

In summary, we elucidated, for the first time, a novel function for FAM76B: modulating inflammation and neuroinflammation by influencing the translocation of hnRNPA2B1. We demonstrated the role of FAM76B in the shared neuroinflammatory pathway of TBI and neurodegeneration, particularly in FTLD-TDP. This study may offer important information for the future development of diagnostic biomarkers and immunomodulatory therapeutics for TBI and neurodegeneration, including for FTLD-TDP.

## Acknowledgement

This work was supported by research grants to H.X. from the National Natural Science Foundation of China (No. 81773265), Key Research and Development Plan of Shaanxi Province (No. 2018SF-106), and the Fundamental Research Funds for the Central Universities (No. GK202007023 and GK202107018).

## Availability of data and materials

The mass spectrometry proteomics data have been deposited to the ProteomeXchange Consortium (http://proteomecentral.proteomexchange.org) via the iProX partner repository [1] with the dataset identifier PXD037539. ([1] Ma J, et al. (2019) iProX: an integrated proteome resource. Nucleic Acids Res, 47, D1211-D1217). All other remaining data are available within the Article and Supplementary Files, or available from the authors upon request.

## Competing interests

The authors declare that no conflict of interest could be perceived as prejudicing the impartiality of the research reported.

## Author contributions

WDY and ZXJ designed this study, performed the experiments and drafted the manuscript. CLH performed the experiments and revised the manuscript. ZJL (Junli Zhao), ZJL (Jiuling Zhu), YPY and LYQ performed the experiments. MQW carried out the data analysis and revised the manuscript. XHB supervised the design of the study and conceived the manuscript. All authors reviewed and approved the final version of this paper. All authors read and approved the final manuscript.

---

**Figure S1.**
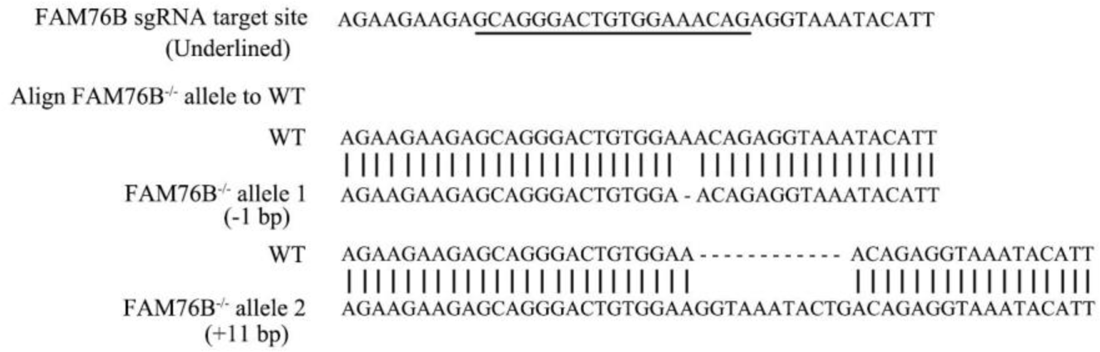
Confirmation by sequencing of Cas9/sgRNA-mediated FAM76B knockout in the U937 cell line. The sequencing of genomic DNA from the *FAM76B^-/-^* U937 cell line showed that one allele contained a 1-bp deletion and the other allele contained an 11-bp addition, as compared to the U937 WT cell line.

**Figure S2.**
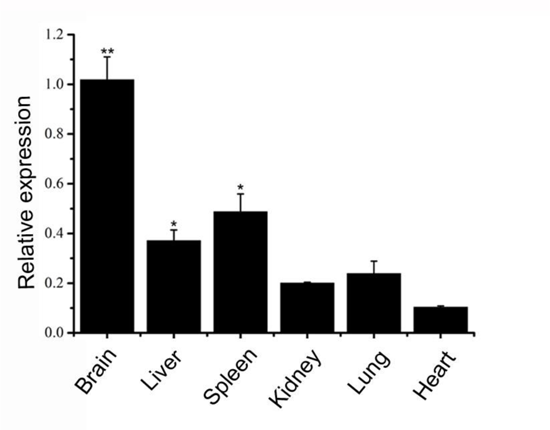
The detection of FAM 76B expression in mouse tissues by real-time PCR. *p<0.05, **p<0.01, statistically significant.

**Figure S3.**
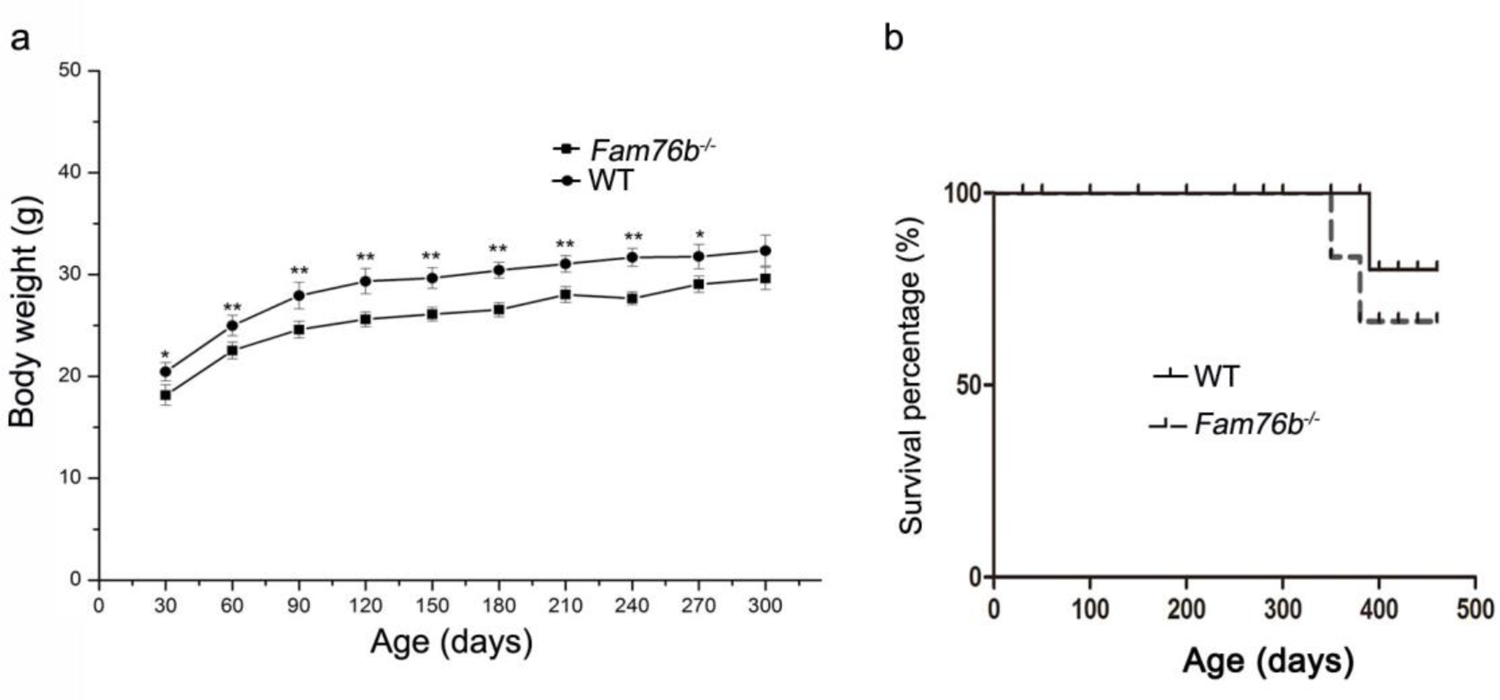
Characterization of *Fam76b* knockout mice. (a) Body weight changes of male WT and homozygous Fam76b mutant (Fam76b^-/-^) mice over time. *p<0.05, **p<0.01, statistically significant. (b) The survival curve of homozygous Fam76b mutant (Fam76b^-/-^) mice did not differ from that of WT mice.

**Figure S4.**
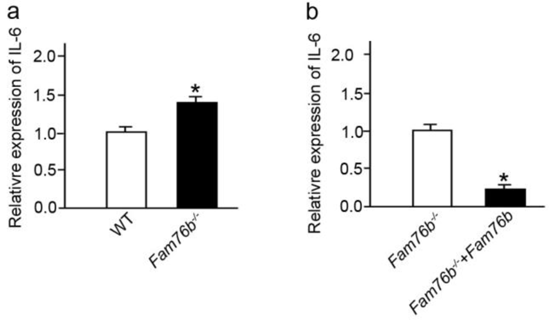
LPS-stimulated bone marrow-derived macrophages from FAM76B knockout mice showed increased IL6 expression. (a) IL-6 expression was revealed by real-time PCR in BMDMs from FAM76B knockout mice in the presence of LPS. (b) IL-6 expression was decreased when the BMDMs from FAM76B knockout mice were infected with FAM76B expressing lentivirus vector in the presence of LPS. The experiments were repeated at least three times. *p<0.05, statistically significant.

**Figure S5.**
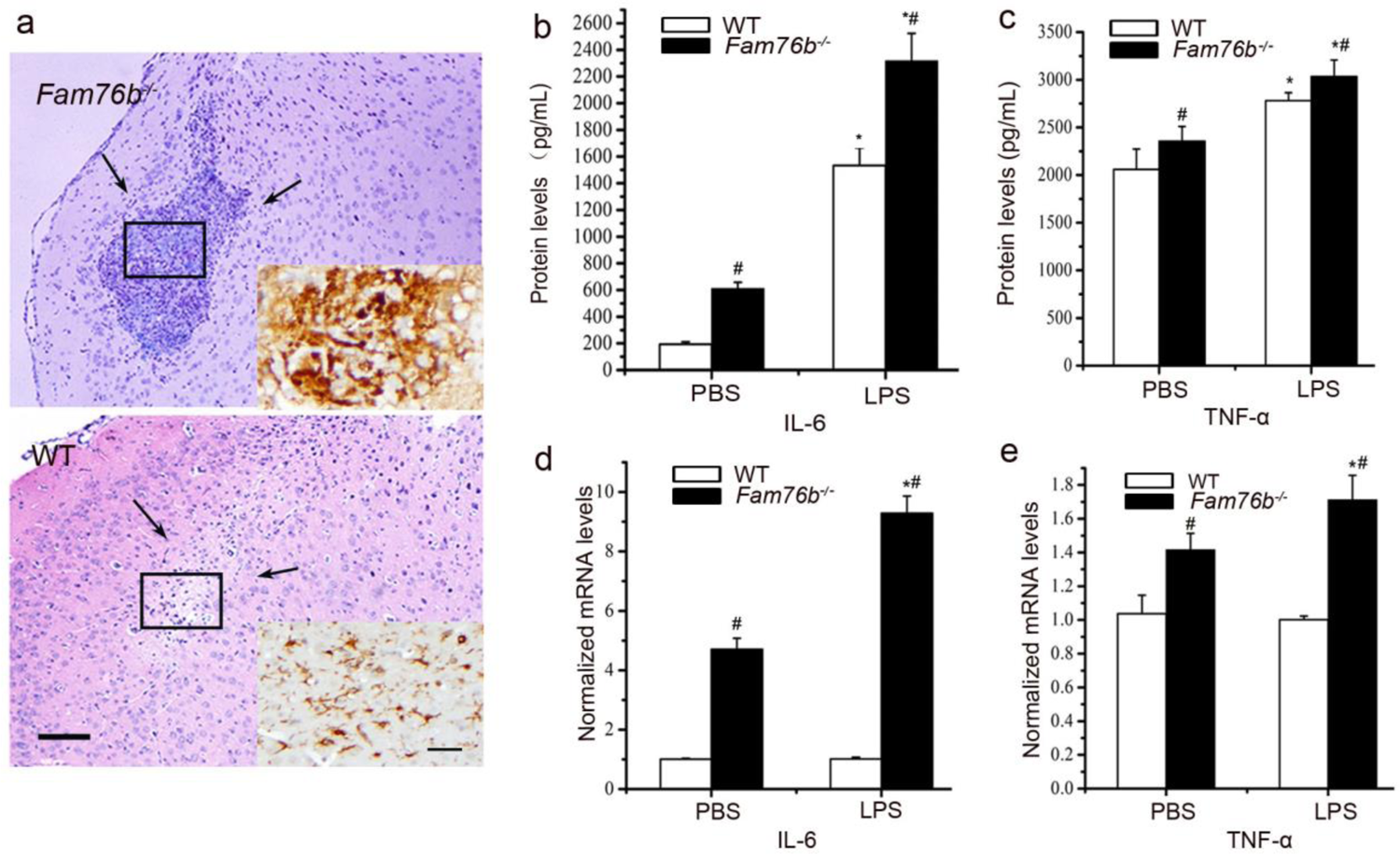
Neuroinflammation in *Fam76b* knockout mice. (a) H&E staining of needle track sites (between the arrows) in LPS-injected mouse brains. A high number of macrophages was observed infiltrating the needle track areas in the LPS-injected brains of *Fam76b* knockout mice. Only a mild inflammatory response was observed at the injection site in WT mice. Scale bar, 100 µm. Insets, the IBA-1-positive cells in needle track sites. Scale bar, 20 µm. (b–e) IL-6 and TNF-α expression at the injection sites of *Fam76b^-/-^* and WT mice post-LPS intracranial injection, by ELISA and real-time PCR. Brain tissues 1 mm in diameter around the needle track were dissected 24 h after the injection and homogenized, followed by ELISA and real-time PCR. The experiments were performed at least three times. *p<0.05, statistically significant, LPS vs. PBS; ^#^p<0.05, statistically significant, *Fam76b^-/-^* vs. WT.

**Figure S6.**
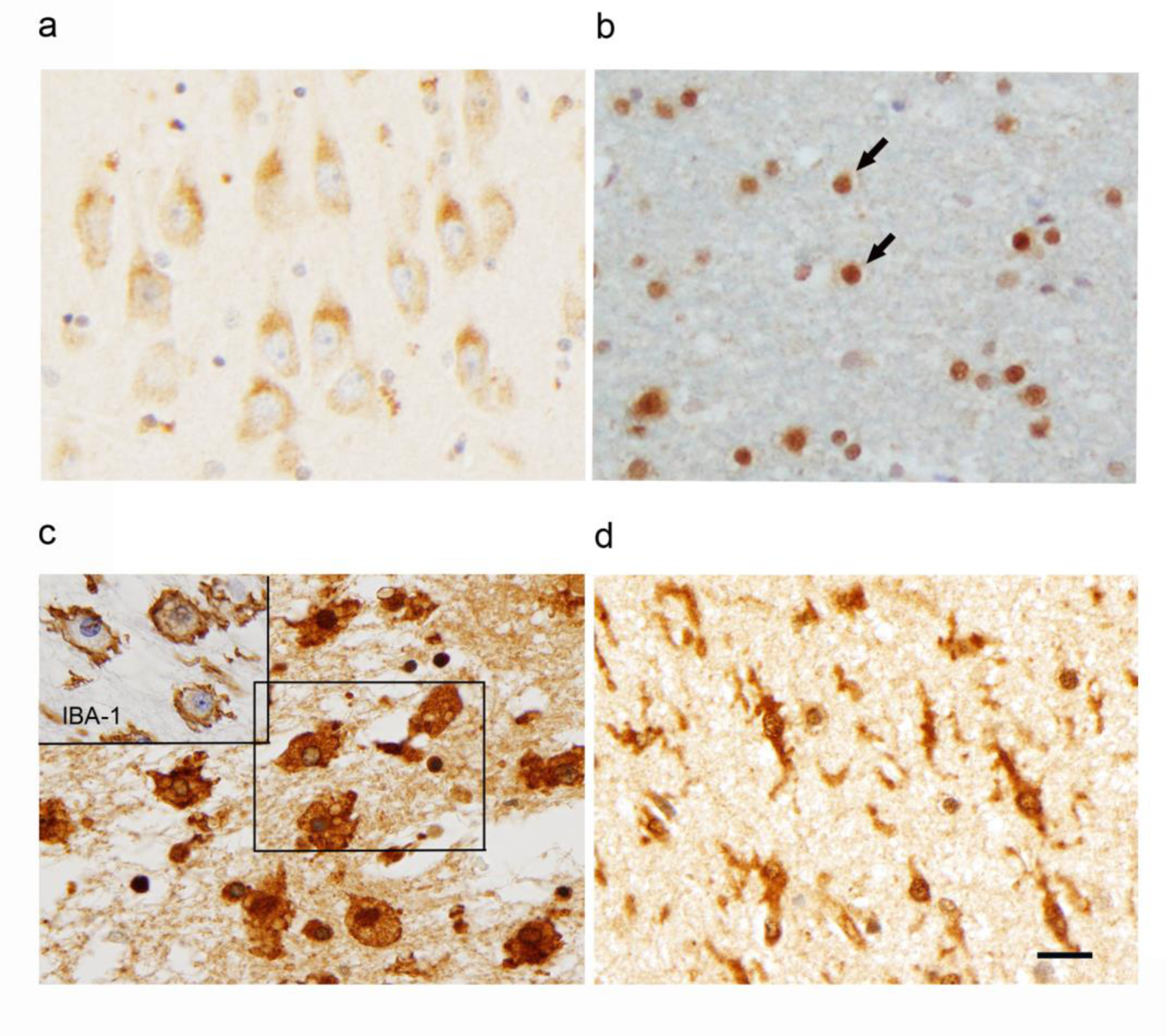
Expression of FAM76B in normal and diseased tissues. The expression of FAM76B in different mouse tissues revealed by immunohistochemical staining demonstrated that FAM76B was localized in the cytoplasm of neurons (a; photo taken of the hippocampal CA1 region) and in the nuclei of glial cells (b; photo taken of white matter). Arrows, oligodendroglial cells of normal human brains. Significant upregulation and cytoplasmic translocation of FAM76B in macrophages in areas of organizing necrosis in the brain of a patient with TBI (c; inset, macrophages labeled by IBA-1 in the mirrored section). Reactive microglial cells in the hippocampus of a patient with acute ischemic injury are strongly immunopositive for FAM76B (d). Scale bar, 20 µm for all photos.

**Table S1.**
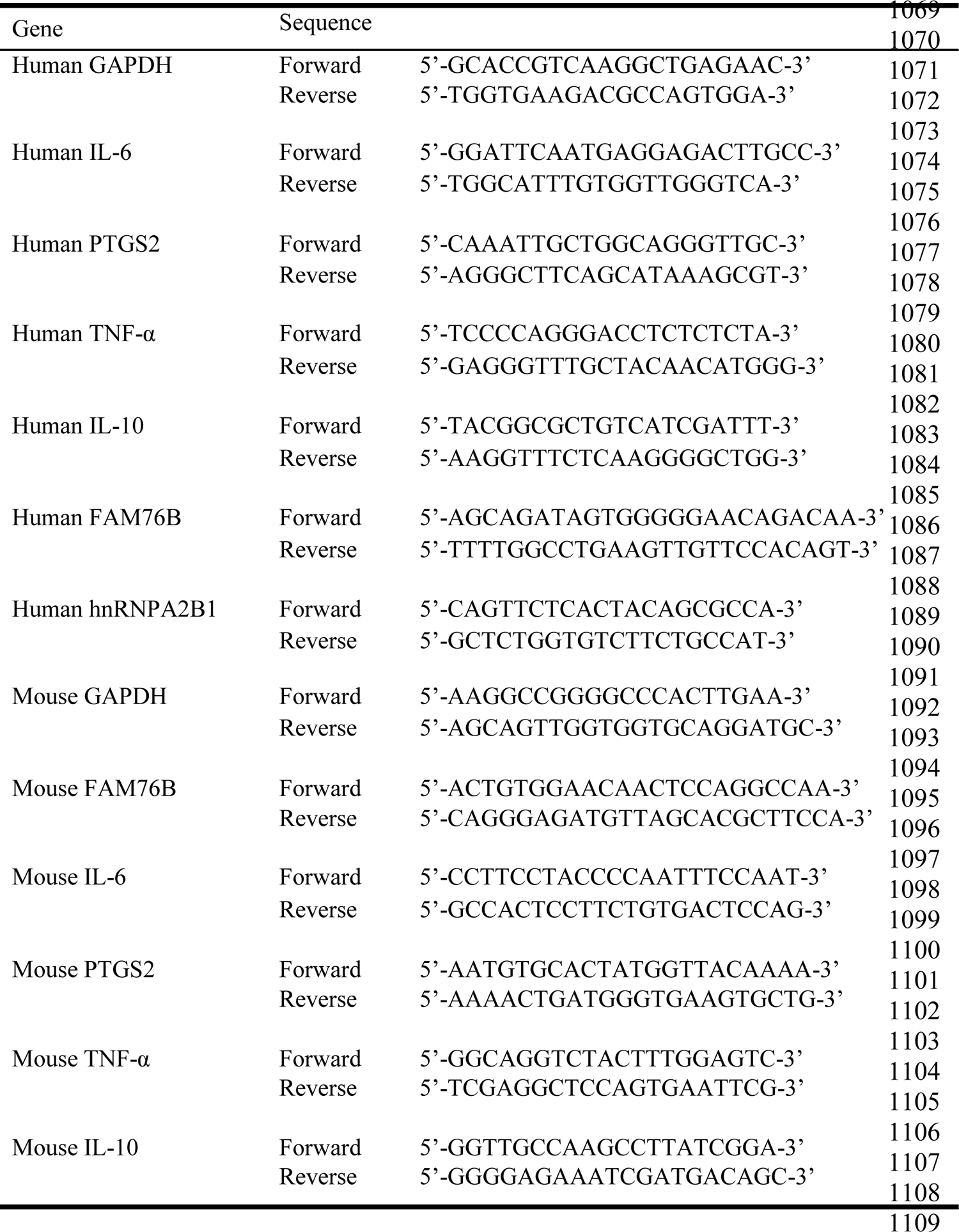
Primers used for real-time PCR

**Table S2.**
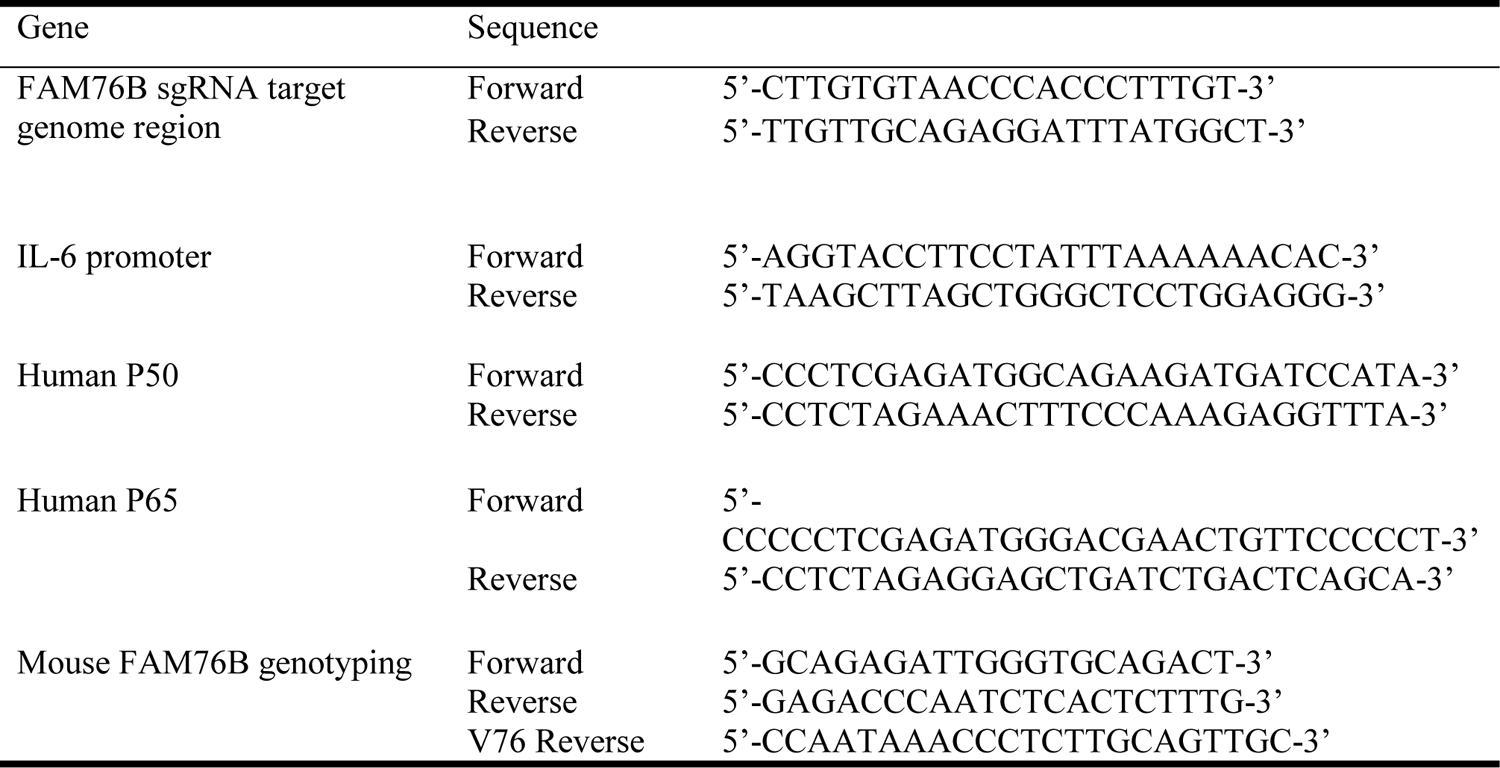
Other primers used in the study

**Table S3.**
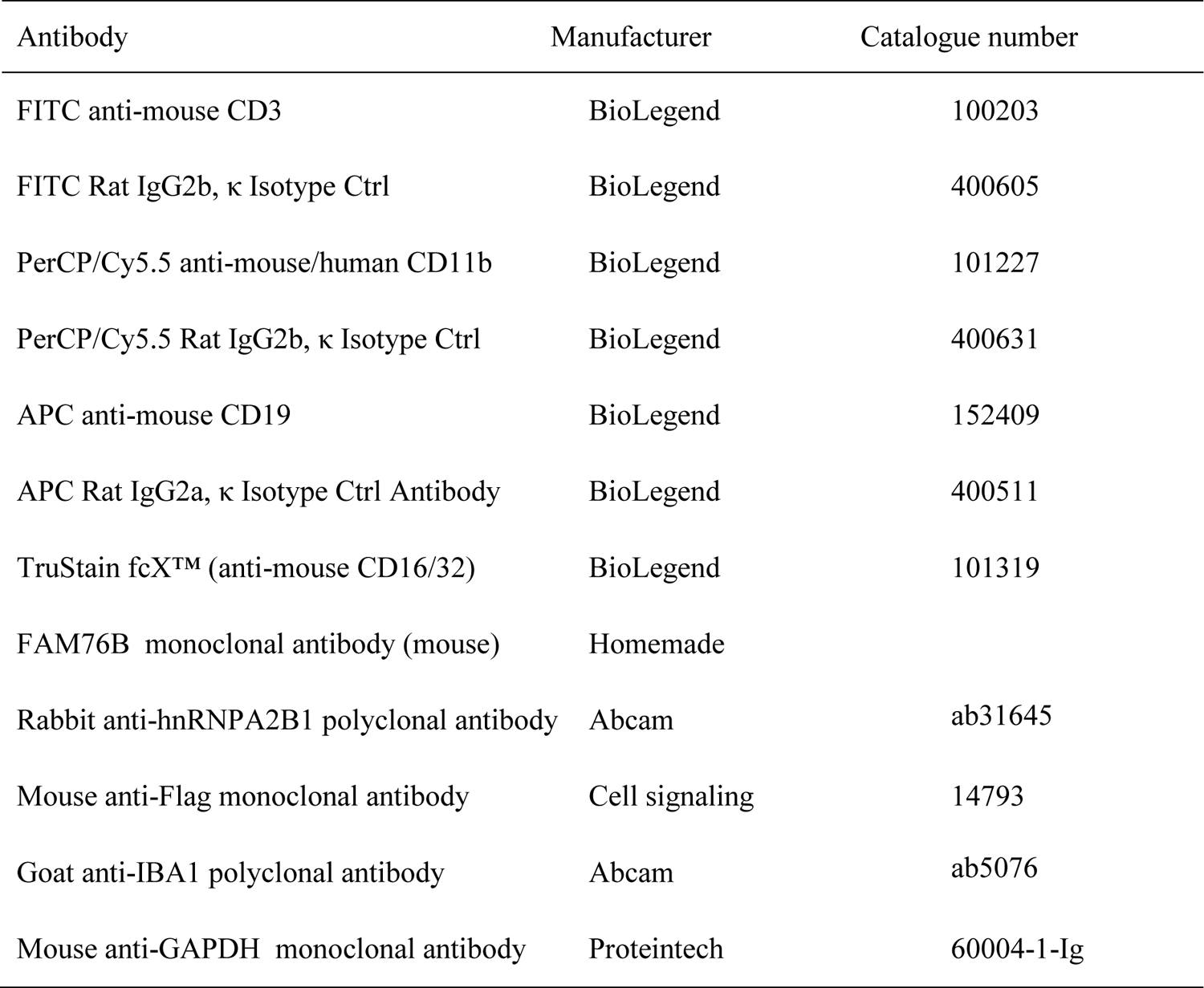
Antibodies used in the study

**Table S4.** Patient demographics

| Neuropathologic diagnosis | Number<br>of cases | Sex<br>M/F | Age at death<br>(mean ± SD) |
| --- | --- | --- | --- |
| CON | 6 | 3/3 | 77.8±7.8 |
| AD | 5 | 2/3 | 77.3±8.4 |
| FTLD-tau | 6 | 2/4 | 72.3±3.2 |
| FTLD-TDP | 6 | 4/2 | 71.5±6.9 |

CON, normal control; AD, Alzheimer’s disease; FTLD-tau, frontotemporal lobar degeneration with tau pathology; FLTD-TDP, frontotemporal lobar degeneration with TAR DNA-binding protein 43 inclusions.

**Table S5.** Proteins that interact with FAM76B and their interacting scores and ranks

| **Protein IDs** | **Protein names** | **Gene names** | **Score** | **Rank** |
| --- | --- | --- | --- | --- |
| **P22626** | Heterogeneous nuclear ribonucleoproteins A2/B1 | HNRNPA2B1 | 164.67 | 5 |
| **P07355** | Annexin A2 | ANXA2 | 60.366 | 11 |
| **P46776** | 60S ribosomal protein L27a | RPL27A | 31.429 | 24 |
| **Q8WWM7** | Ataxin-2-like protein. This gene encodes an ataxin type 2-related protein of unknown function. | ATXN2L | 30.629 | 25 |
| **P09651; Q32P51** | Heterogeneous nuclear ribonucleoprotein A1; Heterogeneous nuclear ribonucleoprotein A1-like 2 | HNRNPA1; HNRNPA1L2 | 27.543 | 28 |
| **P51991** | Heterogeneous nuclear ribonucleoprotein A3 | HNRNPA3 | 27.536 | 29 |
| **P55072** | Transitional endoplasmic reticulum ATPase | VCP | 10.17 | 56 |
| **O75369; P21333; Q14315** | Filamin-B; Filamin-A; Filamin-C | FLNB; FLNA; FLNC | 5.9136 | 99 |
| **P35637; Q92804** | RNA-binding protein FUS | FUS; TAF15 | 5.866 | 101 |
| **P45974** | Ubiquitin carboxyl-terminal hydrolase 5 | USP5 | 5.6403 | 133 |

## Notes

### Competing Interest Statement

The authors have declared no competing interest.

